# The Arabidopsis Class I formin AtFH5 contributes to seedling resistance to salt stress

**DOI:** 10.1101/2024.11.13.623423

**Authors:** Eva Kollárová, Anežka Baquero Forero, Ali Burak Yildiz, Helena Kočová, Viktor Žárský, Fatima Cvrčková

## Abstract

The family of formins, evolutionarily conserved multidomain proteins engaged in the control of actin and microtubule cytoskeleton organization, exhibits considerable diversity in plants. Angiosperms have two formin clades consisting of multiple paralogs, Class I and Class II, the former being often transmembrane proteins located at the plasmalemma or endomembranes. According to available transcriptome data, the *Arabidopsis thaliana* Class I transmembrane formin AtFH5 (At5g54650) exhibits a distinct pattern of transcript abundance in various seedling root tissues with massive increase of transcript level upon salinity stress. To examine a possible role of AtFH5 in NaCl stress response, we generated transgenic plants expressing green fluorescent protein (GFP)-tagged AtFH5 under its native promoter and characterized its tissue and intracellular localization under standard culture conditions and under NaCl stress. While we confirmed the induction of AtFH5 expression by salt treatment, the distribution of tagged protein, with maxima in the border-like cells of the root cap, in the phloem and at lateral root emergence sites, did not reflect previously reported transcript abundance, suggesting posttranscriptional regulation of gene expression. Subcellular localization studies employing also membrane trafficking inhibitors suggested that AtFH5 protein level may be modulated by endocytosis and autophagy. Notably, loss-of-function *atfh5* mutants exhibited increased sensitivity to NaCl stress, indicating that AtFH5 contributes to the development of seedling salt tolerance. These findings highlight the functional importance of AtFH5 in abiotic stress responses.

## 1. Introduction

In response to environmental challenges, plants have evolved sophisticated defence mechanisms to respond and adapt to their environments and ensure their survival (see Zhang et al., 2022). These mechanisms may involve, for instance, epigenetic changes, alterations in gene expression, hormone control, ion channel regulation, redox status and changes in membrane structure and dynamics (Zwiewka et al., 2015; Jiang et al., 2016; Waadt et al., 2022; Kumar and Rani, 2023; Munns and Millar, 2023). Among the diverse abiotic stressors, salinity (NaCl) stress stands out as a prominent challenge for plants. High salt concentrations in the soil result in excessive accumulation of salt in plant tissues, leading to high osmotic pressure, loss of turgor pressure, ionic toxicity, and oxidative stress due to reactive oxygen species (see van Zelm et al., 2020). Additionally, salt stress indirectly impacts plant cell walls by altering the expression of cell wall biosynthesis genes, modifying cell wall enzyme activity, and thereby influencing the physicochemical properties and composition of the cell wall (Dabravolski and Isayenkov, 2023). For example, salt induces pectin deesterification that triggers a cell wall integrity signalling response pathway (Gigli-Bisceglia et al., 2022; see also J. Liu et al., 2021). As a consequence of all these responses, cell volume decreases, cell division slows down, and root and leaf growth are inhibited. Furthermore, salt stress induces stomatal closure, reduces photosynthesis, and limits carbon assimilation (see Chaves et al., 2009). Ultimately, these effects may lead to an early transition from vegetative to reproductive development, premature senescence, and consequently also decreased yield of seed (and possibly other plant parts) with economically relevant consequences in case of crop plants.

The cytoskeleton, composed of microtubules and actin microfilaments, emerges as a pivotal player in the plant’s response to salt stress. Under high salinity, the cytoskeleton undergoes dynamic changes that contribute to perception and transmission of stress signals, thus activating response mechanisms essential for stress adaptation (see Kumar et al., 2022). Microtubules, for example, undergo rapid depolymerization and reassembly in response to salinity stress (Wang et al., 2007). Microtubule-associated proteins and various other factors, promoting either depolymerization or reassembly, exhibit significant gene expression changes upon salt stress (Chun et al., 2021) and contribute to this dynamic behaviour (see, e.g., Wang et al., 2010; Ma and Liu, 2019; Yang et al., 2020; Kumar et al., 2022). These responses are, moreover, intimately linked to various regulatory pathways, such as, e.g., small GTPase signalling (Li et al., 2017) or ethylene response (Dou et al., 2018).

Similarly, the dynamics of microfilaments, encompassing depolymerization and re-polymerization, are vital for plant salt tolerance. In the early stages of salt exposure, high salinity stress initiates the assembly and bundling of microfilaments, whereas prolonged exposure leads to microfilament depolymerization and eventual seedling death. However, stabilizing microfilaments can reverse this detrimental effect. (Wang et al., 2010). Disturbances in actin dynamics also influence reactive oxygen species accumulation under salt stress conditions (Liu et al., 2012). Moreover, microfilament stabilization can compensate for defects in salt stress signalling via the Salt Overly Sensitive (SOS) pathway, likely also involving calcium signalling (Wang et al., 2010; Ye et al., 2013).

Actin-binding proteins (ABPs) are integral in regulating plant salt tolerance. For example, the actin-nucleating ARP2/3 complex contributes to stress response through the modulation of mitochondrial-dependent Ca^2+^ signalling. Arabidopsis mutants with impaired function of the actin nucleating ARP2/3 complex thus exhibit increased salt sensitivity, and this effect appears to be intimately linked with altered regulation of Ca^2+^ homeostasis (Zhao et al., 2013). Another ABP, ADF1, whose expression is controlled by the MYB73 transcription factor, contributes to salt stress response by regulating actin filament organization via its F-actin depolymerizing/severing activity (Wang et al., 2021). Even subtle modulation of microfilament dynamics can affect salinity tolerance. For example, an actin-depolymerizing factor from the halophyte *Spartina alterniflora* confers drought and salt tolerance when constitutively overexpressed in rice better than its rice homolog (Sengupta et al., 2019).

Formins, a family of evolutionarily conserved cytoskeletal organizers, can control actin nucleation, a rate-limiting step in the assembly of actin filaments. They are multidomain proteins containing the conserved Formin Homology 2 (FH2) domain, whose dimer can nucleate and cap actin filaments. This domain is typically preceded by the proline-rich Formin Homology 1 (FH1) domain that interacts with actin-profilin complexes, and a variable N-terminal end. Based on FH2 domain phylogeny and domain organization, angiosperm formins are categorized into two classes: Class I and Class II (Deeks et al., 2002). Class I formins typically have N-terminal transmembrane and extracellular domains, while most Class II formins feature a membrane lipid-binding domain related to the protooncogene PTEN (Phosphatase and Tensin Homolog) at the N-terminal end (Grunt et al., 2008).

Functional studies of plant formins are complicated by the large size of the formin gene family (with 21 paralogs in Arabidopsis) and functional overlap (“redundancy”) between individual paralogs, resulting in rather inconspicuous mutant phenotypes. Nevertheless, formins are emerging as versatile regulators in plants, influencing various cellular processes. They are contributing to the shaping of pavement cells and trichomes (Rosero et al., 2016; Oulehlová et al., 2019; Cifrová et al 2020), to epidermal cell elongation (Cui et al., 2023) and to pollen tube tip growth (Lan et al., 2018; Kollárová et al., 2021), as well as to symplastic transport (Diao et al., 2018). At least some of these functions may be due to transmembrane Class I formins actively orchestrating the interactions between the cytoskeleton and cellular membranes (see Cvrčková et al., 2024).

The Arabidopsis Formin Homologue 5 (AtFH5, At5g54650) is a typical Class I formin with a N-terminal secretory signal, an extracytoplasmic domain and a transmembrane domain preceding the conserved cytoplasmic FH1 and FH2 domains. Its characterization focused so far mainly on its role in the male gametophyte, particularly in comparison with the closely related formin AtFH3, which is predominantly pollen-expressed (e.g. Lan et al., 2018; Liu et al., 2018; Lara-Mondragón et al., 2022, Xu et al., 2024). Compared to its gametophytic function, the role of AtFH5 in the sporophyte remains largely uncharacterized, although publicly available transcriptome data suggest its widespread expression in vegetative tissues. In an earlier overexpression study, AtFH5 primarily localized to the cell plate in roots; its loss of function resulted in delayed endosperm cellularization, indicating its involvement in cytokinesis (Ingouff et al., 2005). Moreover, the AtFH5 gene is subject to epigenetic regulation, potentially complicating transgenic plant studies (Fitz Gerald et al., 2009).

Here we examine the tissue and subcellular localization, as well as the physiological function, of AtFH5. Building on published transcriptome data, we specifically focus on its expression in roots and its function under salt stress conditions. Besides revealing a distinctive tissue-specific expression pattern, we confirmed that AtFH5 expression responds to salt stress, while the subcellular localization of the AtFH5 protein suggested its participation in endosome trafficking and autophagy. Subsequent examination of *fh5* loss-of-function mutant phenotype under salt stress points towards this formińs contribution to salt stress resistance.

## 2. Materials and Methods

### 2.1. Transcriptome data analysis

Initial screen of the AtFH5 transcriptional pattern has been performed using the Arabidopsis eFP Browser (Winter et al., 2007; University of Toronto, 2024). AtFH5-specific data from the following transcriptome datasets, retrieved via the eFP Browser interface, were used for visualization: organ-specific RNA-seq data from Klepikova et al. (2016), root cell type-specific protoplast microarray data from Brady et al. (2007), and stress response microarray data from the AtGenExpress global stress expression data set (Kilian et al., 2007). In case of organ and cell type expression data, values for similar samples (such as individual flowers or adjacent young or old stem segments) were averaged in order to simplify the visual presentation.

### 2.2. Plant materials

The *A. thaliana* T-DNA insertional mutant *fh5-3*, originating from the Col-0 background, was procured from The Nottingham Arabidopsis Stock Center (NASC ID: N664361). Genotyping was performed via PCR using the LP_SALK_152090 and RP_SALK_152090 primers, which were designed using the online T-DNA Primer Design Tool (**Supplementary Materials Table S1**).

The *fh5c* CRISPR mutant in the Col-8 background was created as follows. The single gRNA with no off-target score was generated using the CRISPR-P web tool (Lei et al., 2014). Reverse and forward primers which include single gRNA and BsaI sites, DT1-BsF-FH5 and DT2-BsR-FH5 respectively (for sequence of these and all following primers see **Supplementary Materials Table S1**), were designed by modifying the two-gRNA system described by Xing et al. (2014). The primers were combined and amplified with DreamTaq DNA polymerase (Thermo Fisher Scientific) to create a complete double-stranded DNA, which was subsequently inserted into the pHSE401 vector using Golden Gate cloning (Engler et al., 2008). The resulting construct expressing a single gRNA was transformed into *A. thaliana* Col-8 by the floral dip method employing *Agrobacterium tumefaciens* (Clough and Bent, 1998). Transgenic plants were selected based on hygromycin resistance, and the presence of homozygous single-base insertions was verified by sequencing (used primers FH5seq_F and FH5seq_R). The generated mutant line was then screened for the absence of the Cas9 cassette using Cas9-specific primers, CAS9 fw and CAS9 rv and confirmed by the absence of hygromycin resistance.

### 2.3. Growth conditions

Seeds were surface sterilized for 2 minutes in 70% ethanol, followed by 10 minutes in 50% bleach solution. After three washes, the seeds were placed on a standard medium consisting of ½ Murashige and Skoog (MS) adjusted to pH 5.7, containing 1.6% (w/v) agar, supplemented with 1% (w/v) sucrose unless stated otherwise. The plants were stratified by 1-2 days of post-imbibition storage at 4 °C in the dark and then vertically positioned for cultivation under long-day conditions (16 h light/8 h dark) at 22°C. For liquid substrate cultures, analogous media were used with agar and sucrose omitted.

The *Nicotiana benthamiana* plants were grown on peat pellets (Jiffy, Zwijndrecht, Netherlands) under long-day conditions (16 h light/8 h dark) at 22°C for 2-3 weeks prior to the transient transformations.

### 2.4. Cloning and plant transformation

The vectors for the production of transgenic lines were created using the Gateway cloning system (Invitrogen™) following the manufacturer’s instructions. To amplify the native promoter, encompassing 2300 bp upstream of the AtFH5 (AT5G54650.1) coding sequence, attB-linked oligonucleotide primers, pFH5_rev_GW and pFH5_for_GW, were employed (for sequence of all primers see **Supplementary Materials Table S1**). The AtFH5 genomic sequence, excluding the stop codon, was amplified using primer set AtFH5_for and AtFH5_rev, and the resulting fragment was subsequently amplified with adapter primers AtFH5_full_for_gw and AtFH5_full_rev_gw. The amplified promoter and AtFH5 sequence fragments were introduced into empty vectors, pDONR™ P4-P1R and pDONR™221 (Invitrogen™) respectively, using the BP Clonase II (Invitrogen™). The resulting entry vectors were recombined along with pEN-R2-nGFP-L3 (Boruc et al., 2010) and pB7m34GW,0 (Karimi et al., 2005) using LR Clonase II (Invitrogen™) to create the final expression vector, pAtFH5::FH5-GFP. For the overexpression vector UBQ::AtFH5-GFP, the gateway compatible entry vector carrying, the constitutive ubiquitin 10 promoter, the AtFH5 coding sequence, pEN-R2-nGFP-L3 (Boruc et al., 2010) and destination vector pB7m34GW,0 (Karimi et al., 2005) were recombined. The entry vectors were sequenced using M13 and gFH5sek1, gFH5sek2, gFH5sek3 primers.

The vectors pAtFH5::AtFH5-GFP were introduced into *fh5-3*, *fh5c* and the UBQ::AtFH5-GFP into Col-0 plants by the floral dip method employing *A. tumefaciens* (Clough and Bent, 1998). The positive transformants were selected on the solid medium supplemented with glufosinate-ammonium (20 µg/ml). All lines taken into further analysis exhibited a single-locus transgene, as documented by the transgenés Mendelian segregation (**Supplementary Materials Table S2**).

### 2.5. Co-localization studies in *Nicotiana benthamiana*

For the co-localization studies, we utilized expression vectors containing fluorescent protein markers, specifically ST-RFP (Renna et al., 2005) or p35S::ARA6-mRFP (Ueda et al., 2004) and UBQ:AtFH5-GFP (see **2.4.**), which were introduced into *A. tumefaciens* strain GV3101 through electroporation. *N. benthamiana* leaf infiltration was performed as described previously (Kollárová et al., 2020) using suspensions containing equal amounts of bacteria carrying the marker and the UBQ::AtFH5-GFP corresponding to optical density OD_600_ of 0.08 for each construct, and the viral silencing suppressor p19 at OD_600_ of 0.05. We examined the fluorescent proteins 48 hours after infiltration using confocal microscopy.

### 2.6. Esculin staining

For phloem staining with esculin (Knox, 2019), a 2.5% solution of the adjuvant Velocity® was applied to both cotyledons of 8 days old seedlings for 1h prior to the esculin loading. A stock solution of esculin (Merck CAS 531-75-9) was prepared by dissolving 20 mg in 1 ml of 70% ethanol and heating for 5 minutes at 95 °C. The stock solution was then diluted to a final concentration of 5 mg/ml working solution, and 2 μl of this solution was applied to both cotyledons. Plant observations were conducted 20 minutes after esculin application.

### 2.7. FM4-64 staining

4-5-day-old seedlings were incubated in a solution of ½ MS medium supplemented with 2 μM FM4-64 dye for 5 to 30 minutes and subsequently examined at defined time points using a confocal microscope.

### 2.8. Drug treatments

The experimental procedures were carried out using 5 days old seedlings, using a ½ MS liquid medium. In the case of the Brefeldin A (BFA) treatment, seedlings were immersed in a medium supplemented with 50 μM BFA and 1 μM FM4-64 for 50 minutes before subsequent observation. For the Wortmannin (WM) treatment, the seedlings were transferred to a medium supplemented with 33 μM WM or DMSO (10,000 x diluted) as a control for 2 hours prior to 5 minutes of FM4-64 staining immediately before observation. For TOR pathway inhibition treatments, seedlings were transferred to a medium supplemented with DMSO (10,000 x diluted), 1 μM AZD8055 (AZD), 1 μM Concanamycin A (ConcA), or a combination thereof for 6 hours before they were observed.

### 2.9. Abiotic stress treatments for protein expression analysis

For the sucrose treatment, seeds were sown on ½ MS solid medium, both with and without 1% sucrose. After 5 days of cultivation, confocal microscopy images were acquired for the purpose of fluorescence quantification.

For applying salt and mannitol stress, seedlings were initially cultivated for 4-5 days on a standard growth medium. Subsequently, they were transferred to the same standard medium but supplemented with either 150 mM NaCl, or 300 mM mannitol or to control media without osmotic supplements, and allowed to culture for a period of 20-24 hours. The imaging for fluorescence quantification was conducted by SDCM.

### 2.10. Imaging

The microscopic images were acquired using either confocal laser scanning microscopy (CLSM) or spinning disc confocal microscopy (SDCM) as specified in the image descriptions. CLSM was performed using a Zeiss LSM880 microscope with Plan-Apochromat 20 ×/0.8 objective or a C-Apochromat 40×/1.2 W Korr FCS M27 objective. The fluorophore was excited with the 488 nm argon laser (GFP) or 561 nm laser and detected by a sensitive 32-channel Gallium arsenide phosphide (GaAsP) spectral detector. The SDCM was performed using a Zeiss Axio Observer 7 microscope with a vertical stage equipped with a Yokogawa CSU-W1 spinning disk unit, alpha Plan-Apochromat 100×/1.46 Oil immersion objective, laser lines set at 488 nm or 561 nm and the PRIME-95B Back-Illuminated Scientific CMOS Camera.

Images for phenotypic observations were captured utilizing the open-source MULTISCAN phenotyping platform built according to Slovak et al (2014). The gel image was obtained using an Azure 600 (Azure Biosystems) gel documentation system.

All images were processed using the Fiji software (Schindelin et al., 2012), applying adjustments for brightness, contrast, and artificial colouring while preserving the fidelity of the original images (Cvrčková, 2019). Three-dimensional images were generated using MorphographX software (Barbier de Reuille et al., 2015).

### 2.11. Fluorescence quantification

Fluorescence was measured using Fiji software, with settings for area integrated intensity and mean grey value. Subsequently, the measured values were converted into corrected total cell fluorescence (CTCF) by subtracting the value of the selected area (corresponding to a single cell unless stated otherwise) multiplied by the mean fluorescence of background readings from the integrated density value (Fitzpatrick, 2014).

### 2.12. Salt and mannitol treatments and phenotype analysis

Five-day-old seedlings initially grown on standard ½ MS medium were transferred to media supplemented with the indicated combinations of 250 mM NaCl, 10 mM CaCl2 or 300 mM mannitol or to control media without osmotic supplements and cultivated for an additional five days. Seedlings cultured on the control NaCl-free medium underwent daily scanning for 5 days to determine the effects of transfer on root growth, followed by root length measurement using Fiji software. The seedlings exposed to salt treatment were evaluated to ascertain the survival rate, with cotyledon bleaching serving as an indicator of seedling death.

### 2.13. Protein isolation and Western blot

Proteins were extracted from 1 g of fresh, whole 2-weeks-old seedlings (treated as indicated in 3.3 and then removed from media and dried) by first pulverizing the tissue in liquid nitrogen. The powdered tissue was then mixed with 1 ml of lysis buffer per gram of tissue (150 mM NaCl, 0.1% Triton X, and 50 mM TRIS HCl-pH 8, supplemented with 13:1 of Complete Mini, Roche).

Samples were vortexed, incubated on ice for 1 hour, and centrifuged (15 minutes at 4 °C, 3620 g). The supernatant was transferred to a new tube and centrifuged again (15 minutes at 4 °C, 16000 g). The final supernatant was used for analysis. Protein concentration was measured using the Bradford assay (Bio-Rad) and normalized to 100 μg for pAtFH5:AtFH5 and 5 μg for 35S samples to account for expression level differences.

The samples were mixed 5:1 with SDS loading buffer and loaded on 10% polyacrylamide gel (ran 300 V/40 mA, 1.5 h). The transfer onto nitrocellulose membrane was done using the Trans-Blot Turbo Transfer System (Bio-Rad) and Trans-Blot Turbo RTA Mini 0.2 µm Nitrocellulose Transfer Kit (Bio-Rad) according to the manufacturer’s protocol. The predefined MIXED MW protocol was used (1.3 A, 25 V for 7 minutes). The membrane was blocked with agitating overnight at 4 °C in PBS with 5% milk and 0.05% Tween, followed by washing 3 times in PBS with 0.05% Tween for 5 mins.

The membrane was incubated for 100 minutes with the primary antibody (affinity purified polyclonal rabbit anti-GFP, Agrisera AS15 2987, stock concentration 1 μg/μl, diluted 1:3000 in 3% milk in PBS), followed by three PBS washes. The secondary antibody (Goat anti-rabbit IgG HRP, Promega, diluted 1:133333 in 5% milk in PBS) was then applied for 50 minutes, followed by three PBS washes. After the final wash, the membrane was split to separately expose the pAtFH5:AtFH5-GFP and free GFP samples due to differences in signal intensity. Visualization was performed using Amersham ECL Prime Detection Reagent (Cytiva) following the manufacturer’s instructions. The membrane was exposed for 3 minutes and visualized on the Azure 600 (Azure Biosystems).

### 2.14. Statistics and data visualization

Statistical analyses were conducted employing One-way ANOVA with a post-hoc Tukey HSD (Honestly Significant Difference) test carried out using an online R-based calculator (Vasavada, 2016). Boxplots were generated using the BoxPlotR tool (RRID: SCR_015629; Spitzer et al., 2014) and charts were created in Microsoft Excel. Chi square tests have been performed using an online calculator (Stangroom, 2024).

## 3. Results

### 3.1. Tissue localization of AtFH5 expression in seedling roots

Analysis of publicly available transcriptome data indicated high levels of AtFH5 transcript in roots, in particular at the seedling stage, with maxima in the root cap, stele and mature trichoblasts (**Figure 1 A, B**). RNA level in the root appeared to be massively increased upon prolonged NaCl treatment (**Figure 1 C**). In order to experimentally ascertain the tissue localization of AtFH5 in *A. thaliana* plants and for further study of the subcellular localization of this protein, we generated constructs encoding AtFH5 C-terminally tagged with the green fluorescent protein (GFP) under the control of either its native promoter (pAtFH5) or the constitutive UBQ promoter. Subsequently, we introduced the pAtFH5::AtFH5-GFP construct into the *fh5-3* T-DNA insertion mutant, as well as into the *fh5c* mutant, generated using the CRISPR-Cas9 technique (**Supplementary Materials Figure S1**). In parallel, we also introduced the overexpressing UBQ::AtFH5-GFP construct into wild-type plants.

**Figure 1.**
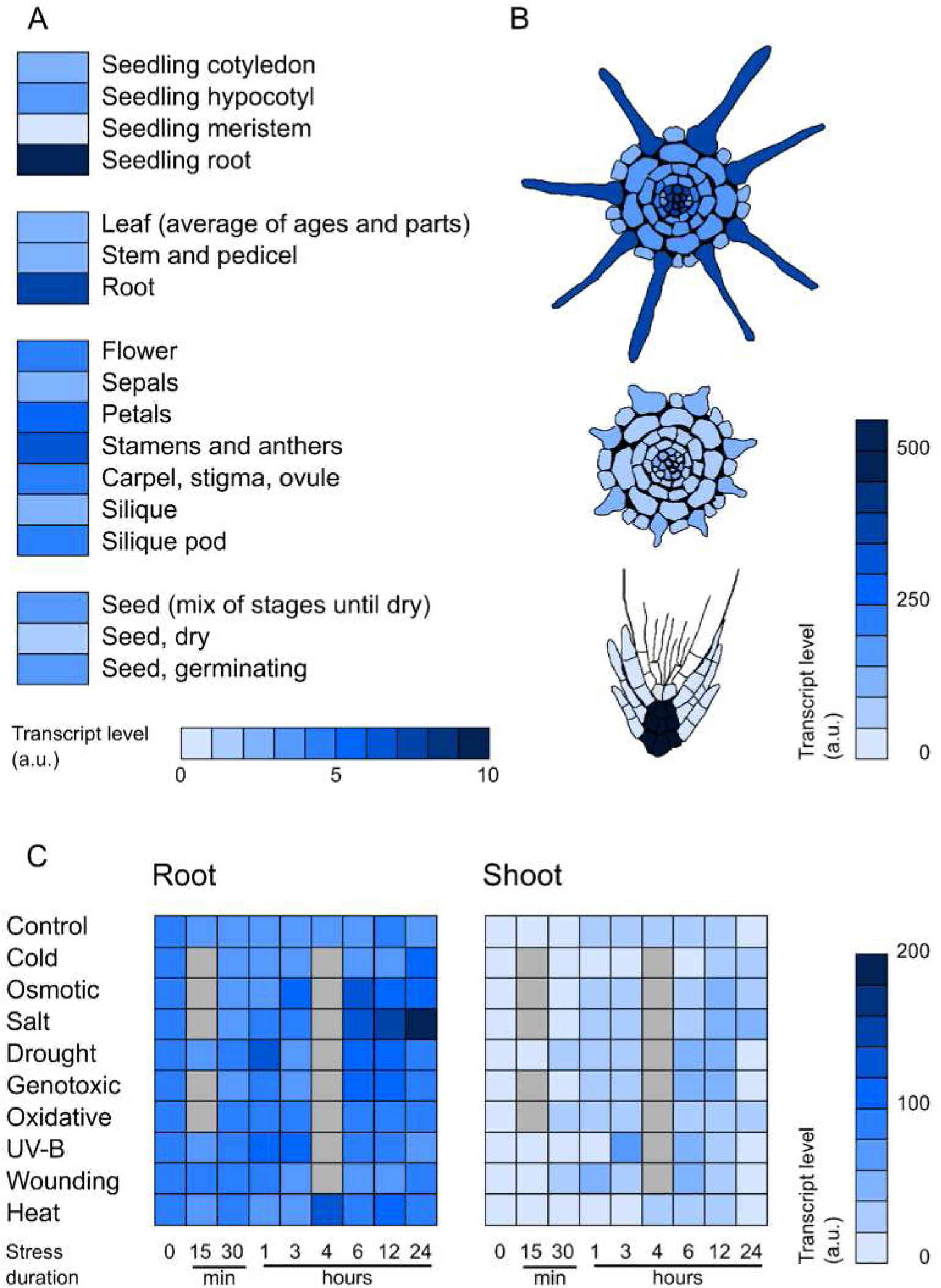
Expression profile of AtFH5 as inferred from public transcriptome data. (A) Transcript abundance in selected organs. (B) Tissue-specific transcript levels in the developing root. Top: mature root, middle: young differentiation zone, bottom – columella, root cap and quiescent centre. (C) Response of AtFH5 transcript levels to selected abiotic stresses. All heatmaps show transcript abundance on a linear scale in arbitrary units.

Using confocal microscopy, we investigated the tissue-specific expression of AtFH5-GFP under native promoter in roots of transgenic seedlings. We observed a relatively strong fluorescent signal in the root cap (**Figure 2 A**), especially in the gradually detaching layers of border-like cells (Vicré et al., 2005). Remarkably, while the signal was detected in the root cap layers fully or partially attached to the root, it was absent in the cells that had already undergone separation from the root tip (**Figure 2 B, C**).

**Figure 2.**
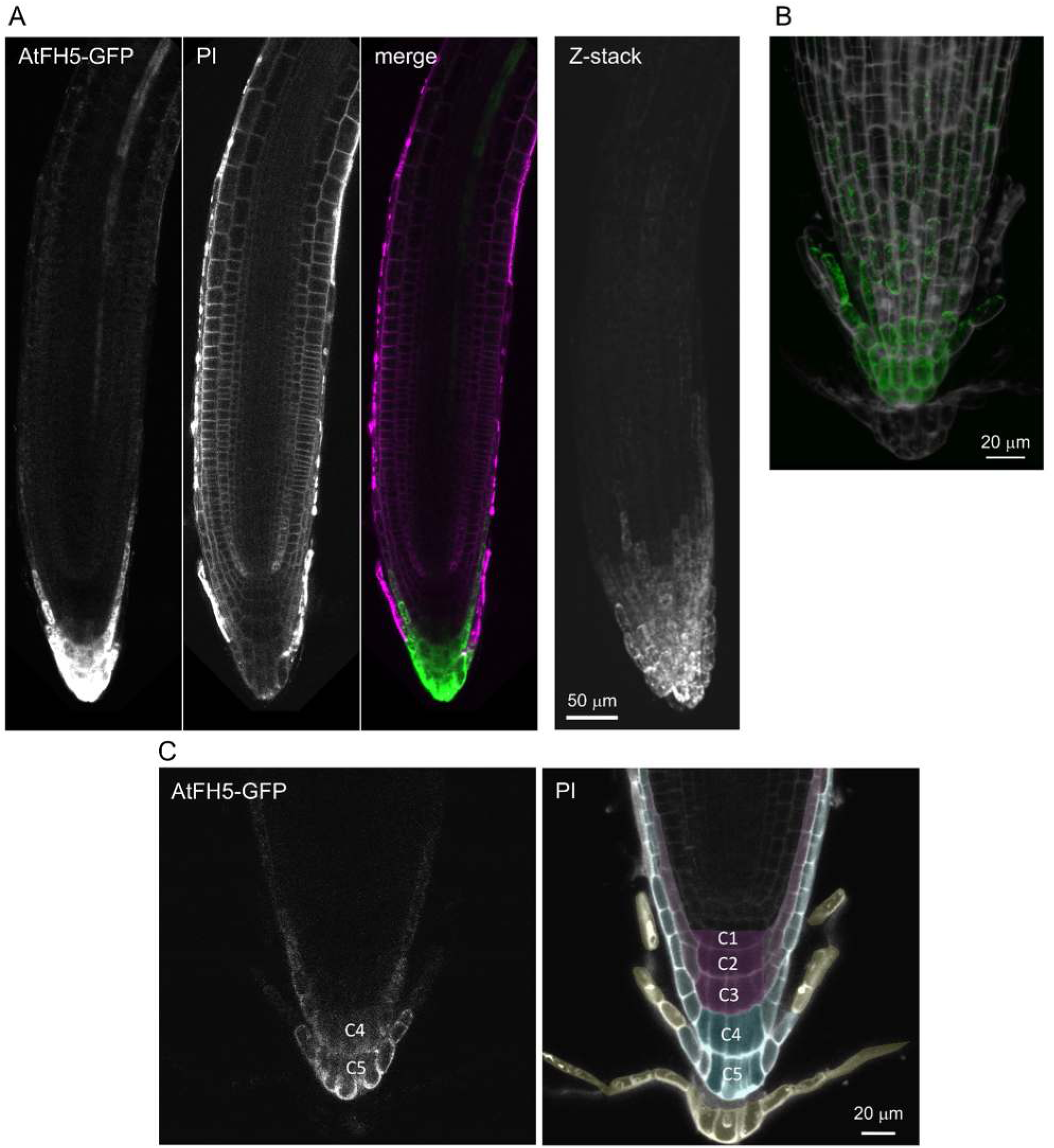
Tissue localization of pAtFH5::AtFH5-GFP expression in the seedling root tip. (A) Left: Longitudinal CLSM optical sections of a root stained by propidium iodide (PI). Single channel images are shown in grayscale, the merged display shows GFP in green and PI in magenta. Right: Z-projection of a CLSM Z-stack of the same plant. (B) 3D reconstruction of a CLSM Z-stack of PI stained root cap. PI in grayscale, GFP in green. (C) Longitudinal CLSM optical sections of a root tip stained by PI. Left: GFP, right: PI, yellow – separated root cap cell layers, cyan – root cap cell layers with GFP signal, magenta – root cap cell layers without GFP signal, C1 – C5: columella layers.

Consistent with the transcriptome data, high expression was also observed in the stele in a pattern suggesting localization in the phloem and its companion cells (**Figure 3 A**). To verify that the protein is localized within the phloem, we stained the transgenic plants with the fluorescent coumarin glucoside esculin, which is efficiently loaded into the phloem via the sucrose transporter, rendering it a suitable tracer for labelling phloem (Knoblauch et al., 2015). As expected, esculin demonstrated co-localization with AtFH5-GFP (**Figure 3 B**).

**Figure 3.**
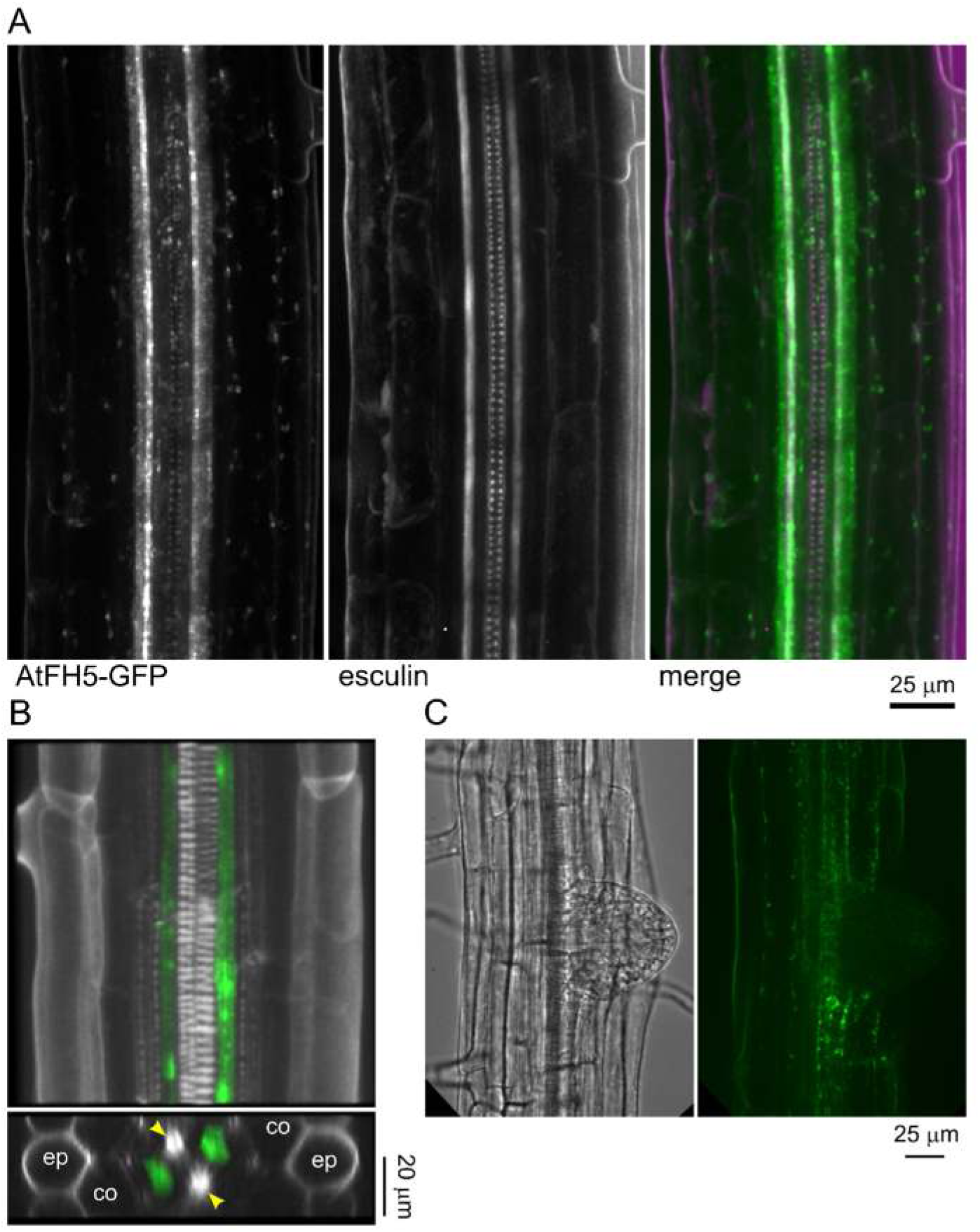
Tissue localization of pAtFH5::AtFH5-GFP in the root stele and in an emerging lateral root. (A) Longitudinal SDCM optical sections of a seedling root stained by esculin. Single channel images are shown in grayscale, the merged display shows GFP in green and esculin in magenta. (B) Top: 3D reconstruction of a CLSM Z-stack of a PI stained root. Bottom: Transversal optical section of the same root. PI in grayscale, GFP in green. Yellow arrowheads indicate the xylem pole; ep, epidermis; co, cortex. (C) Maximum intensity projections of a CLSM Z-stack of a freshly emerged lateral root primordium. Left – brightfield, right – GFP.

While we did not detect the trichoblast signal expected from the transcriptome data, AtFH5-GFP fluorescence was also observed in rhizodermal, and possibly also cortical, cells separating to provide space for lateral root emergence (**Figure 3 C**).

An analogous expression pattern was also seen in the *fh5c* background, whereas the overexpressing line displayed a signal all over the root (**Supplementary Materials Figure S2 A, B**). In dividing cells of the late meristematic zone of the overexpressing plants AtFH5-GFP was found in the cell plate (**Supplementary Materials Figure S2 C**), consistent with a previous report (Ingouff et al., 2005).

### 3.2. AtFH5-GFP localizes to endomembrane compartments in border-like cells

At the subcellular level, AtFH5-GFP formed motile puncta in the cytoplasm of root cap cells (**Figure 4 A**, **Supplementary Materials Movie S1**). To examine the relationship of these puncta to compartments of the endomembrane system, we incubated the seedlings with the FM4-64 styryl dye that is internalized through endocytosis and sequentially labels plasmalemma, early endosomal, late endosomal, and vacuolar compartments (Rigal et al., 2015). Since the AtFH5-GFP-containing compartments co-localized with internalized FM4-64 within mere 15 minutes of incubation, and some FM4-64 signal overlapping with that of AtFH5 was seen already after five minutes (**Figure 4 B, C**; **Supplementary Materials Movie S2**), AtFH5-GFP appears to localize to early endosomes.

**Figure 4.**
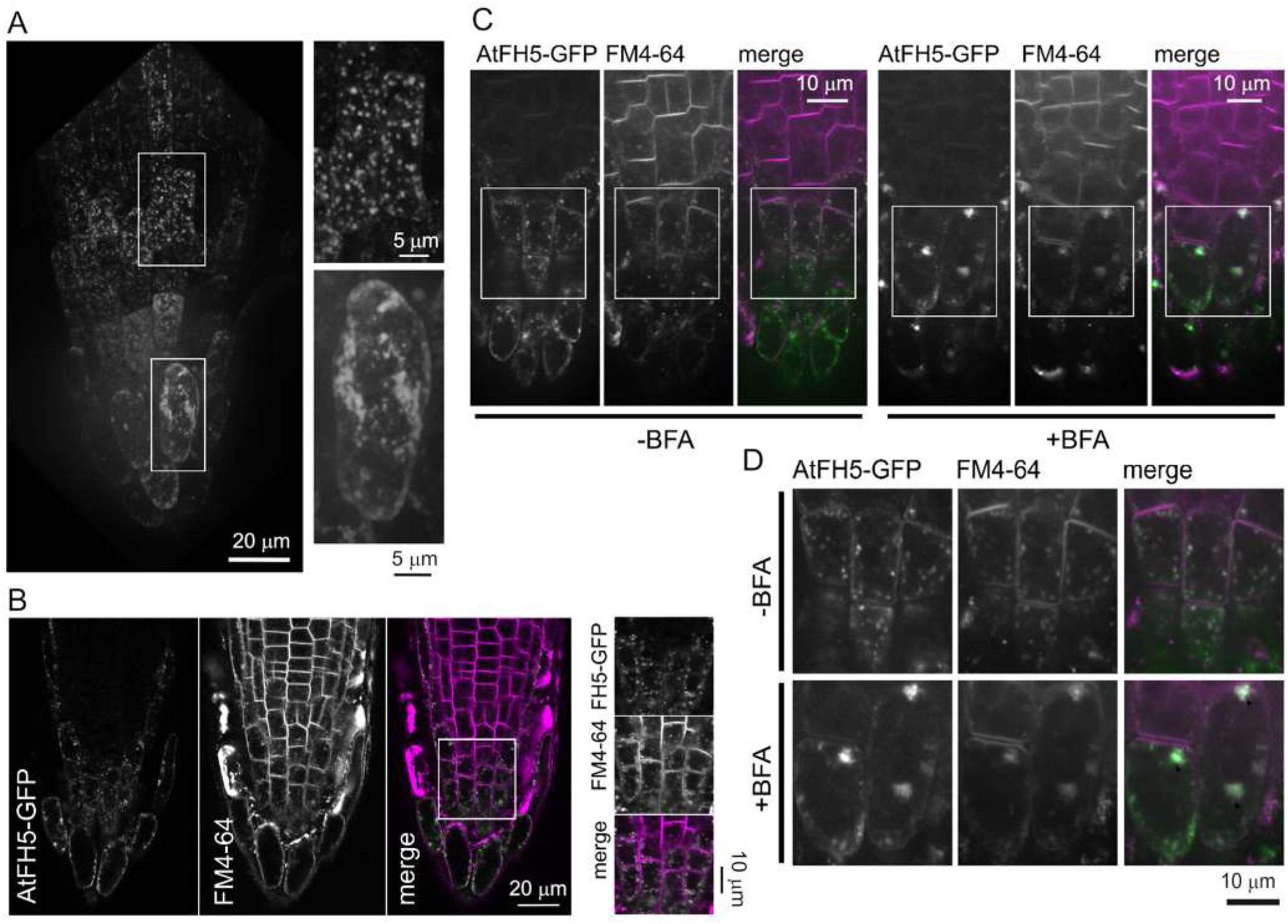
Subcellular localization of AtFH5-GFP in the root cap. (A) Left: 3D reconstruction of a SDCM Z-stack of a seedling root cap. Right: Close-up images of the marked areas. (B) Left: Longitudinal SDCM optical sections of a root cap stained for 5 min by FM4-64. Single channel images are shown in grayscale, the merged display shows GFP in green and FM4-64 in magenta. Right: Close-up images of the marked area. (C) Longitudinal SDCM optical sections of a root cap stained by FM4-64 after 50 min BFA treatment with mock control. Single channel images are shown in grayscale, the two channels in the merged display are represented by green (GFP) and magenta (FM4-64). (D) Close-up images of the marked area from (C). All images are from pAtFH5::AtFH5-GFP seedlings.

Plant endosome trafficking can be targeted by the fungal toxin brefeldin A (BFA) that inhibits specific ADP ribosylation factor/guanine nucleotide exchange factors (ARF-GEFs), causing the endocytic tracer FM4-64 to aggregate rapidly within vesicle aggregates referred to as BFA compartments (Geldner et al., 2003; Ritzenthaler et al., 2002). To determine whether the AtFH5-GFP-marked endosomes are BFA-sensitive, transgenic plants were treated with BFA. As shown in **Figure 4 D**, the AtFH5-GFP signal relocated to BFA-induced bodies. Additionally, employing transient heterologous co-expression of AtFH5-GFP and the Golgi marker ST-RFP in *Nicotiana benthamiana* epidermal cells, we observed close association (but not overlap) of the AtFH5-GFP-labeled bodies with Golgi apparatus itself indicating TGN/EE association of AtFH5 (**Supplementary Materials Figure S3**, **Movie S3**).

### 3.3. AtFH5-GFP is directed to the vacuole via the autophagy pathway

To further elucidate the character of AtFH5-GFP-containing compartments, we examined the effect of wortmannin (WM), an inhibitor of the biosynthesis of the signalling phospholipids phosphatidylinositol 3- and 4-phosphates (Wang et al., 2009). Application of WM induces the late endosomes to dilate or form ring-shaped structures, but does not noticeably affect the Golgi apparatus and early endosomes (Jaillais et al., 2008; Tse et al., 2004). Following the WM treatment, increased accumulation of AtFH5-GFP was observed in the root cap (**Figure 5 A, B**; **Supplementary Materials Movie S4**), and increased AtFH5-GFP abundance after WM treatment was confirmed also by Western blots (**Figure 5 C**). Intriguingly, in WM-treated plants, the AtFH5-GFP signal was predominantly located at the plasmalemma and cell cortex and exhibited a diffuse distribution, while in the control it showed a characteristic punctate pattern, concentrating at compartments labelled with FM4-64 (**Figure 5 D**). Transient co-expression of the late endosome marker ARA6-RFP (Jaillais et., 2008; Ueda et al. 2004) with AtFH5-GFP in *N. benthamiana* leaf epidermis shows AtFH5-GFP partly localizing to the late endosome, supporting the late endosomal identity of some of the AtFH5-GFP labelled compartments (**Supplementary Materials Figure S4, Movie S5**), in addition to the above-reported TGN/EE localization of this formin.

**Figure 5.**
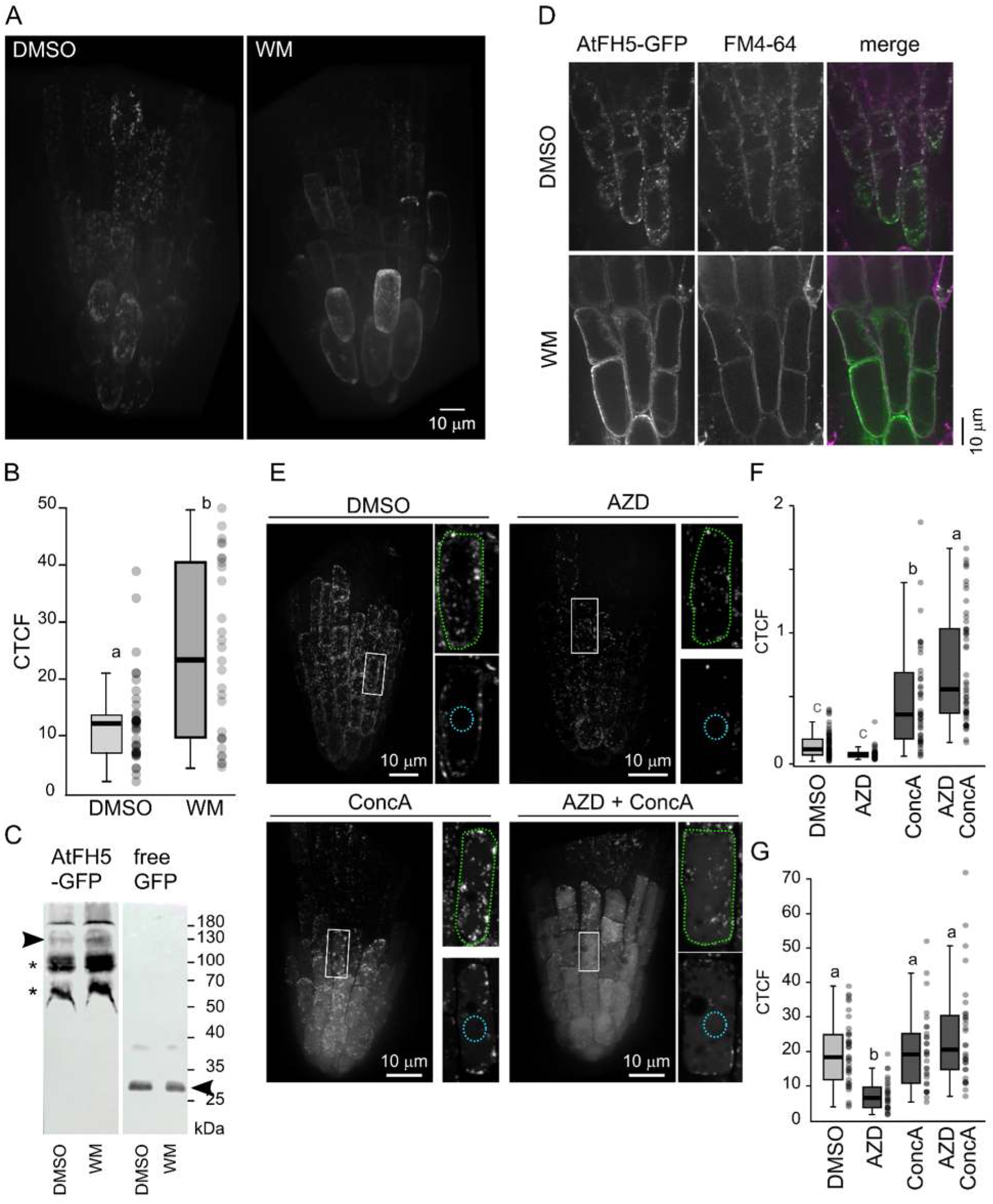
AtFH5-GFP localizes to TGN and late endosomes in the root cap. (A) 3D reconstruction of SDCM Z-stacks of AtFH5-GFP seedling root caps treated with DMSO (control) or WM. (B) Corrected total cell fluorescence (CTCF) of AtFH5-GFP signal in border-like cells treated with DMSO or WM (arbitrary units). (C) Western blot of proteins from 10 days old transgenic seedlings treated with DMSO or WM. Identical amount of total protein was loaded for both samples from each genotype; however, there was more protein in AtFH5-GFP than in free GFP controls, resulting in some nonspecific background (asterisks). Arrows: predicted position of AtFH5-GFP and free GFP. (D) SDCM optical sections of AtFH5-GFP seedlings stained by FM4-64 after control or WM treatment. Single channel images in grayscale, GFP in green and FM4-64 in magenta in the merged display. (E) Maximum projections of SDCM Z-stacks of AtFH5-GFP seedling root cap treated with DMSO or the indicated drugs. Insets: top – maximal projection of the marked area; bottom – a single optical section from the centre of the same cell; outlines denote areas quantified in (F) and (G). (F) CTCF of AtFH5-GFP signal of a standard-sized region within the vacuole (cyan outline); arbitrary units. (G) CTCF of AtFH5-GFP signal of a whole cell section (green outline); arbitrary units. Treatment duration was 2 h in (A) to (D), 8 h in (E) to (G). Letters in (B), (F) and (G) denote statistical significance of the observed differences (one-way ANOVA, p < 0.05).

To further document that AtFH5-GFP is directed to the vacuole via autophagy related pathway, we applied the Target of Rapamycin (TOR) signalling pathway inhibitor AZD8055 (AZD), which induces autophagy in plant cells, either as a sole treatment or in combination with Concanamycin A (ConcA) which inhibits V-ATPase activity, preventing vacuolar acidification that would result in quenching of any GFP fluorescence present in the vacuole, and causing preservation of autophagic cargo that was delivered to the vacuole (see Klionsky et al., 2021). When measuring fluorescence in the central portion of an optical scan of a single cell, corresponding to the cell’s vacuole, we did not observe any differences between control and AZD-treated roots. However, upon simultaneous ConcA application, increased vacuolar fluorescence was observed in AZD-treated plants, indicating signal accumulation in the vacuole (**Figure 5 E, F**). When measuring fluorescence throughout the entire cell (marked in Figure XD as a green area), we observed a decrease in overall signal for samples treated with AZD compared to controls. Conversely, when ConcA was applied, no such difference was found, mirroring the signal in DMSO controls without additional treatment (**Figure 5 G**). Moreover, seedlings grown for 5 days on a sucrose-containing medium, i.e. under conditions stimulating the TOR pathway (Dai et al., 2022) and thus suppressing autophagy, exhibited a significant increase in cytoplasmic fluorescence compared to controls without sucrose (**Supplementary Materials Figure S5**). These observations support the notion that AtFH5-GFP is directed to the vacuole through an autophagic process.

### 3.4. AtFH5 expression is modulated by salt and osmotic stress

According to transcriptomic data, the AtFH5 RNA level in roots is increased upon prolonged 150mM NaCl treatment (**Figure 1 C**). To confirm the expression pattern inferred from public transcriptome data, expression of AtFH5-GFP was followed in roots of pAtFH5:AtFH5-GFP seedlings treated with 150 mM NaCl for 24h (**Figure 6 A, B, C**). Besides the root cap and phloem signal that was also observed under standard conditions, subtle fluorescence was also detected in the epidermal cells. Upon higher magnification, this signal exhibited the above-described characteristic particulate structure (**Figure 6 B**). A statistically significant increase of overall fluorescence intensity was observed in the root elongation zone (**Figure 6 D**), but not in the root cap (**Figure 6 E**).

**Figure 6.**
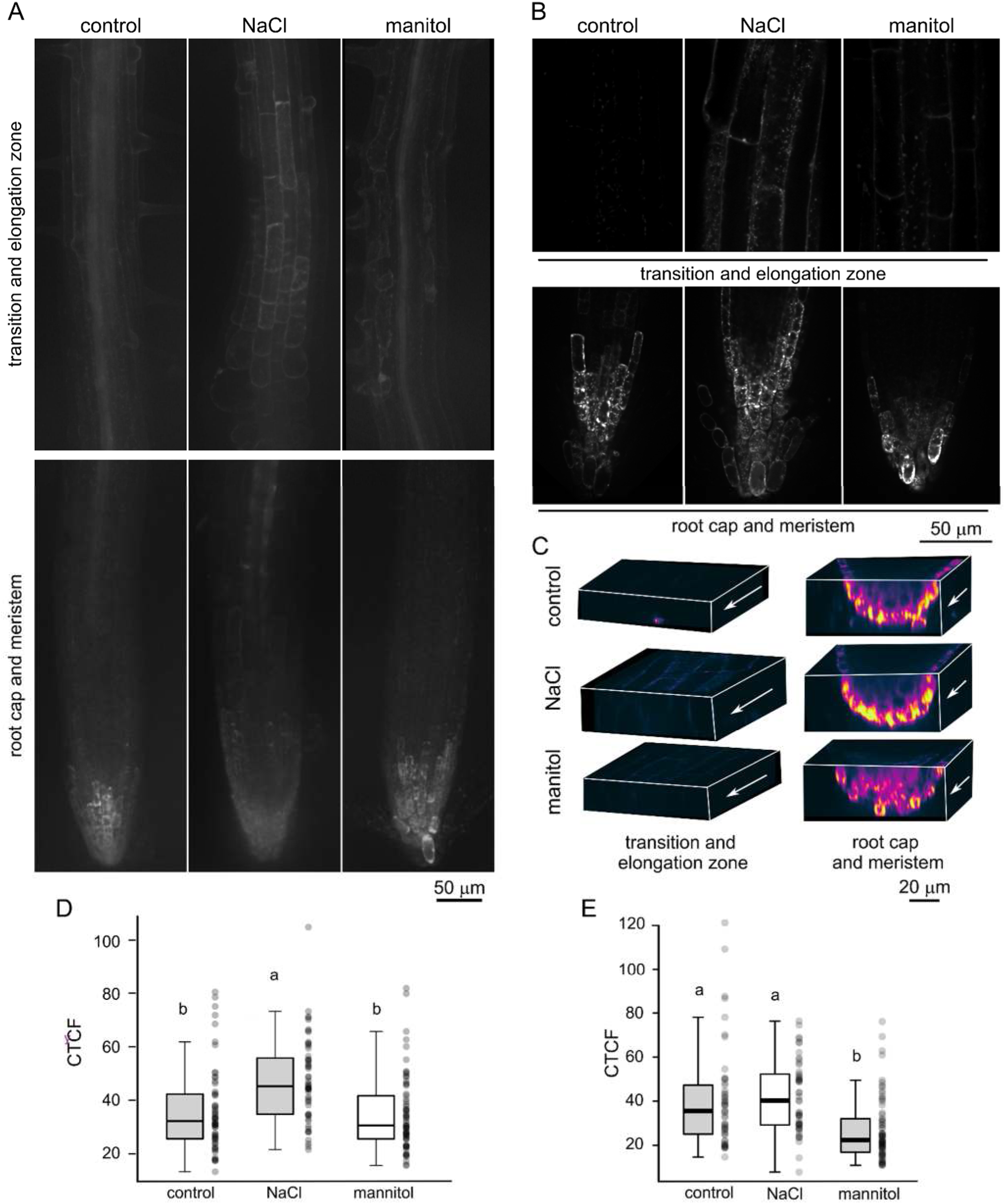
Effects of salt and osmotic stress on pAtFH5::AtFH5-GFP expression. (A) Maximal projections of SDCM Z-stacks from the indicated zones of seedling roots incubated for 24 h under control conditions or in media containing 150 mM NaCl or 300 mM mannitol. (B) Single SDCM optical sections from the indicated zones of seedling roots incubated as in (A). (C) 3D reconstructions from the indicated zones of seedling roots incubated as in (A); artificial colouring using the Warm Helix colour map, arrows denote the rootward direction. (D) CTCF quantification of AtFH5-GFP signal in a standardized cell area in the root cap from seedlings treated as in (A). (E) CTCF quantification of AtFH5-GFP signal in a standardized cell area in the root elongation zone from seedlings treated as in (A). Letters in (D) and (E) denote statistical significance of the observed differences (one-way ANOVA, p < 0.05).

Since transcriptomic data indicate increased AtFH5 expression also upon exposure to high mannitol concentrations, suggesting a possible role of this formin in generalized osmotic stress response, we also examined the effects of mannitol treatment on AtFH5-GFP expression. Unlike salt treatment, exposure to 300mM mannitol did not cause obvious changes in AtFH5-GFP fluorescence compared to control seedlings (**Figure 6 A, B, C**), and no significant change in fluorescence intensity was found in the elongation zone epidermis (**Figure 6 D**). The fluorescence level in the root cap was, surprisingly, significantly decreased in mannitol-treated plants compared to untreated controls (**Figure 6 E**).

### 3.5. AtFH5 contributes to salt stress resistance

Since the expression pattern of AtFH5 suggests that it may participate in salt stress response, we investigated the phenotypes of loss of function mutants *fh5-3* and *fh5c*, as well as AtFH5-GFP complemented and overexpressing transgenic plants, under high salinity conditions. Using cotyledon bleaching as an indicator of seedling damage or death, we examined seedling survival 5 days after transfer on a high-salinity (250 mM) medium (**Figure 7 A**). The transfer to a fresh agar plate and associated mechanical damage per se did not affect seedling survival or growth in any of the genotypes examined (**Figure 7 B**). After five days on high-salinity medium, both mutant lines exhibited a significantly higher fraction of damaged or dead seedlings compared to wild type and AtFH5-GFP overexpressing plants, as well as to complemented *fh5-3* plants expressing AtFH5-GFP from its native promoter (**Figure 7 C**). This indicates that AtFH5 contributes to salt stress resistance and that our AtFH5-GFP fusion protein is biologically functional, since it complements the increased salt sensitivity observed in loss-of-function mutants.

**Figure 7.**
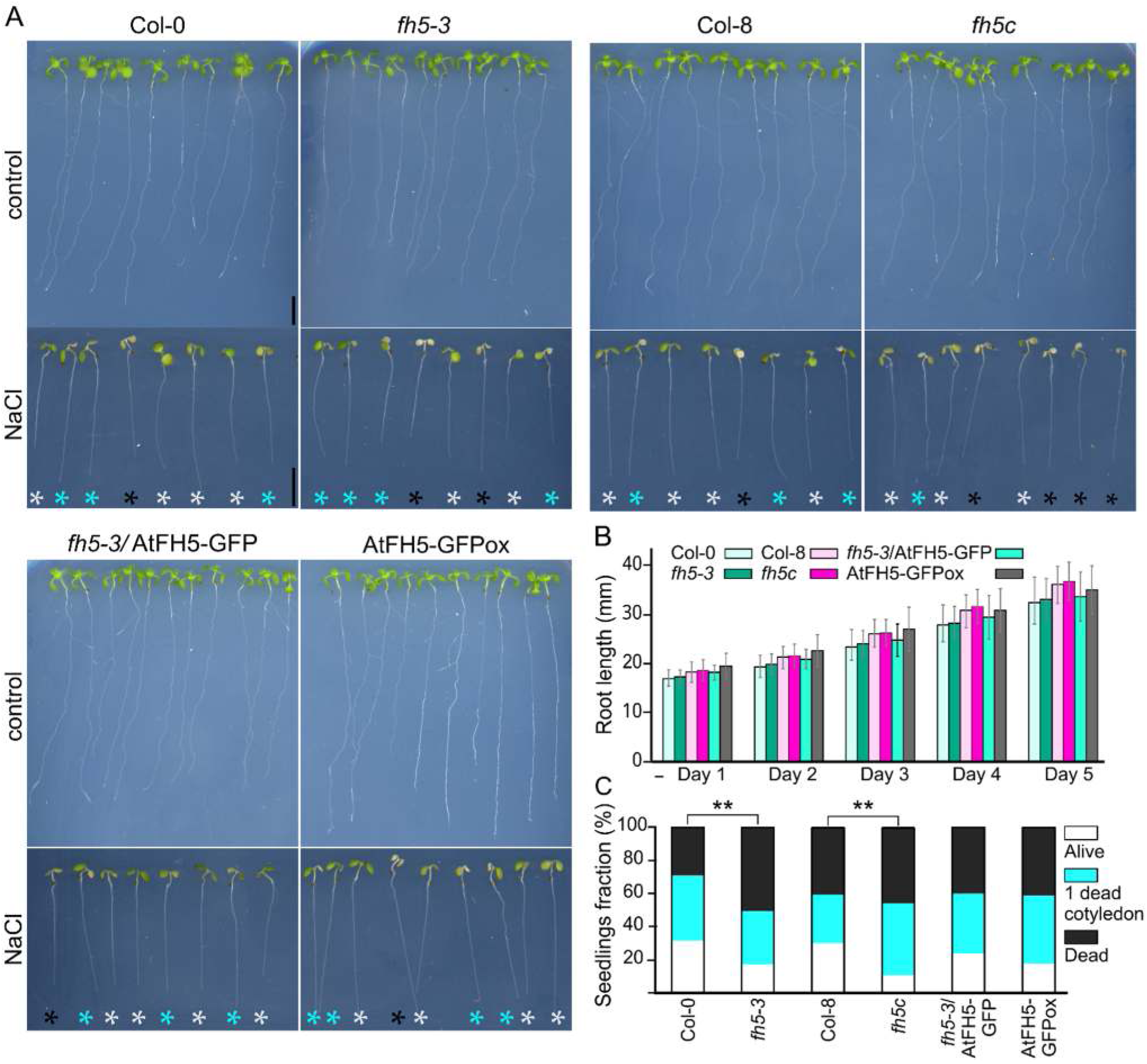
Loss of AtFH5 results in increased seedling damage under salt stress. (A) Representative seedlings after 5 days of exposure to 0 and 250 mM NaCl. Black asterisks – bleached seedlings, cyan asterisks – seedlings with one bleached cotyledon, light grey asterisks – surviving plants. (B) Primary root length of seedlings of the indicated genotypes during five days after transfer to a new standard medium plate, showing that any observed inhibition or damage is not caused by seedling transfer. Between-genotype differences are not significant at any time point (one-way ANOVA, p > 0.05). Data presented are means ± SD, n ≥ 30 per genotype. (C) Fraction of seedlings exhibiting various extent of damage after 5 days on media with 250 mM NaCl. Black – bleached seedlings, cyan – one bleached cotyledon, white – surviving seedlings. Asterisks denote significant differences (p < 0.01).

The increased sensitivity of *atfh*5 mutant seedlings towards NaCl is not a mere consequence of impaired response towards osmotic stress, because mannitol treatment, although inhibiting root growth to a comparable extent in all genotypes examined (**Supplementary Materials Figure S6**), does not cause seedling lethality. Moreover, in all AtFH5 genotype observed, the NaCl toxicity is alleviated by simultaneous CaCl_2_ treatment, which also stimulates root growth when applied on its own but does not counteract the inhibitory effect of NaCl (**Supplementary Materials Figure S7**). This agrees with previous reports indicating that Ca^2+^ can contribute to the resistance to a variety of abiotic stresses and is specifically involved in reaction to NaCl toxicity (Wilkins et al., 2016; Steinhorst et al., 2022). We therefore conclude that AtFH5 contributes specifically to seedling resistance towards salt stress.

## 4. Discussion

In this study, we are examining the expression, intracellular localization and possible stress response-related role of the Class I formin AtFH5, an actin nucleator whose transcriptional response to salt treatment suggests a possible role in salinity response. AtFH5 has been so far studied mainly with focus on its role in cytokinesis (Ingouff et al., 2005) and in the male gametophyte, where it localizes to the plasmalemma and endomembrane compartments at the tips of emerging or elongating pollen tubes and cooperates with the closely related formin AtFH3 in highly polarized vesicle trafficking during pollen germination and pollen tube elongation (Lan et al., 2018; Liu et al., 2018; Ruan et al., 2023; Lara-Mondragón et al., 2022). We are exploring its role in roots, where transcriptomic data reveal a relatively high expression, particularly in the rhizodermis, stele, and columella. Our transgenic plants expressing a biologically active AtFH5-GFP fusion protein under its native promoter did not exactly copy the transcriptome pattern, exhibiting instead signal maxima in the root cap, especially in border-like cells, as well as in the phloem and in cells adjacent to emerging lateral roots, suggesting post-transcriptional regulation of AtFH5 expression. An overexpressing line involving the UBQ promoter, we observed expression throughout the entire root, consistent with constitutive activity of this promoter.

In the root meristem, AtFH5 was reported to accumulate at the cell plate, a membrane-rich structure formed through highly polarized vesicular trafficking (Ingouff et al., 2005). Some other Class I formins, especially AtFH1, AtFH4 and AtFH8, exhibit enrichment at cell plates or transverse cell-to-cell contacts on the plasma membrane (Deeks et al., 2005; Oulehlová et al., 2019), and many also localize to the plasmodesmata. However, this has not been demonstrated for AtFH5, which showed a uniform distribution across the plasma membrane instead (Diao et al., 2018, Oulehlová et al., 2019). Our overexpressed AtFH5-GFP decorated the cell plate in dividing cells, consistent with previous findings (Ingouff et al., 2005). In agreement with transcriptome data, AtFH5-GFP expressed from its native promoter was not detected in the root meristematic zone, at least under standard culture conditions. However, the protein was found in border-like cells of the root cap, where it displayed a weak uniform distribution on the plasma membrane and an intense motile punctate signal in the cytoplasm. Additionally, differences in punctate signal distribution in the cytoplasm were noted between the outermost layer and the second outermost layer at the root cap periphery, likely due to endomembrane compartments rearrangement during maturation and detachment of border-like cells (Goh et al., 2022). Similar to another Class I formin, AtFH1 (Oulehlová et al. 2019), the punctate AtFH5-GFP signal co-localized with both early and late endosomes, as documented by the kinetics of FM4-64 staining and wortmannin sensitivity of these compartments. Following BFA treatment, AtFH5-GFP accumulated within BFA bodies; similar co-localization with endosomes was observed in male gametophytes (Liu et al., 2018). Heterologous transient expression experiments suggested that AtFH5-GFP-containing compartments associate with the Golgi apparatus without directly overlapping with it, and may thus correspond to the trans-Golgi network. Thus AtFH5, similar to other transmembrane formins, relocates between secretory pathway compartments, the plasmalemma and endosomes. Ultimately, it is directed into the vacuole, as documented by pharmacological experiments employing wortmannin, a phosphatidylinositol 3-kinase and phosphatidylinositol 4-kinase inhibitor known to affect protein vacuolar sorting and endocytosis (Takáč et al., 2012).

To confirm that AtFH5’s vacuolar targeting is related to autophagy, we employed an inhibitor of Target of rapamycin (TOR) to stimulate autophagic processes, resulting in relocation of AtFH5-GFP from the cytoplasm to the vacuole, supporting the hypothesis that AtFH5 is directed to the vacuole through an autophagic pathway. Interestingly, recent studies indicate that reduced ATP levels, which result from suppression of TORC1 activity (or mitochondrial dysfunction), cause actin filaments to become less dynamic and less responsive to LatB-induced depolymerization, although the overall organization of actin remains unaffected. A possible explanation for the observed reduction in actin dynamics could involve an increased ratio of ADP-G-actin to ATP-G-actin, which is known to slow actin polymerization (Dai et al., 2022). Alternatively, changes in actin-binding protein activity may also contribute to this effect.

AtFH5 has been demonstrated to play a critical part in regulating the dynamics of actin microfilaments in germinating pollen grains. Specifically, AtFH5 localizes to vesicles and precedes actin filament assembly, with profilin proteins like PRF4 and PRF5 enhancing its interaction with actin filaments, potentially promoting its processivity (Liu et al., 2018; C. Liu et al., 2021). Additionally, the increase in AtFH5-GFP levels upon sucrose addition to the growth medium supports a possible contribution of energy availability to the regulation of AtFH5 level. It is tempting to speculate that AtFH5 may be directed to the vacuoles via autophagy-related pathways under energy-deficient conditions as a part of a cellular strategy to regulate actin dynamics.

In seedlings grown under standard conditions, we observed a high level of AtFH5-GFP signal in border-like cells of the root cap, while on NaCl-containing media, we found an increase in AtFH5-GFP fluorescence intensity in the transition and elongation zones rhizodermis, while no notable expression changes were detected in the root cap. Interestingly, a root-cap-specific proteomics study did not identify AtFH5 among the most abundant proteins in this tissue under control or salt stress conditions (Das et al., 2023). However some wheat (*Triticum aestivum)* formins are upregulated in response to high salinity, including TaFH1 and TaFH9, close relatives of AtFH5 (Duan et al,. 2021). Furthermore, a comparative proteomic analysis of *Prunella vulgaris* seedlings revealed a significant upregulation of Profilin 3, an actin-binding protein that interacts with the FH1 domain of formins, in response to salt treatment (Z. Liu et al., 2021).

Salt stress is known to exert distinct short-term and long-term effects on plant growth. In the short term, it induces osmotic stress by limiting water availability, while in the long term, it leads to ion toxicity due to salt accumulation, disrupting cellular ion homeostasis (see Munns, 2005). We thus also examined the effects of osmotic stress imposed by mannitol treatment, to confirm that the upregulation of AtFH5 is not related to osmotic stress alone. Indeed, we found no increase in AtFH5 levels in the rhizodermis of seedlings exposed to high concentrations of mannitol, and even observed decreased signal in the root cap, possibly linked to osmotic stress-induced autophagy, which may lead to the targeting of AtFH5 to the vacuole in the root cap upon autophagy induction.

While the precise mechanisms remain unclear, it is evident that the depolymerization of microfilaments significantly impairs a plant’s ability to endure salt stress, whereas their polymerization is crucial for enhancing salt tolerance (Wang et al., 2009). Our observation that loss of function mutants of the AtFH5 formin, which has been shown to participate in the regulation of microfilament dynamics, exhibited impaired survival on high NaCl media, and this phenotype was effectively rescued upon introduction of AtFH5-GFP, indicates functional importance of AtFH5 in mitigating salt stress. Additionally, our findings reveal that treatment with CaCl₂ rescues salt toxicity, consistent with previous studies indicating that exogenous calcium ions (Ca²⁺) can reduce sodium ion (Na⁺) influx in plants, as observed in both *Arabidopsis* (Shabala et al., 2006) and rice (Roy et al., 2019).

Our protein localization observations indicate that AtFH5 is a cargo of secretory and autophagic trafficking. Interestingly, autophagy is induced by salt stress in Arabidopsis and contributes to adaptation to salinity (Luo et al., 2017). The decreased resistance of AtFH5 mutants to NaCl stress suggests that this formin actively participates in cell-level salt stress resistance by a mechanism that may involve the function of the autophagic pathway under high salinity conditions. If this is the case, AtFH5 may act as an active cargo of membrane trafficking (sensu Cvrčková et al., 2024) – i.e., a protein embedded in the membrane of autophagy-related compartments and modulating their fate while underway towards its final vacuolar destination.

## 5. Conclusion

We have identified the Arabidopsis class I formin AtFH5 as a new player in the salt stress response. Building on a combination of physiological, pharmacological and cytological observations, we hypothesize that this formin may be participating to the interplay of cytoskeletal rearrangements, exocytosis and autophagic membrane trafficking in the primary root tissues, contributing thereby to salt stress adaptation in young seedlings.

## CRediT authorship contribution statement

E.K. - Conceptualization, Visualization, Investigation, Formal analysis, Writing – original draft; A.B.F. - Conceptualization, Investigation, Formal analysis, Writing – review and editing; A.B.Y. - Investigation, Validation; H.K. - Investigation, Writing – review and editing; V.Ž. - Conceptualization. Writing – review and editing; Funding acquisition, F.C. - Conceptualization, Visualization, Writing – review and editing; Supervision, Funding acquisition.

## Acknowledgements

We thank Katarína Kulichová for the Gateway compatible UBQ10 promoter construct, Radek Bezvoda for assistance using the Multiscan platform, Zdeněk Erben from ADAMA CZ for providing the sample of Velocity**®,** Ondřej Šebesta for expert imaging assistance, and Marta Čadyová for technical support.

## Appendix.#Supplementary materials

Supplementary figures, tables and video files descriptions (PDF)

Supplementary Movie S1 (MP4)

Supplementary Movie S2 (MP4)

Supplementary Movie S3 (MP4)

Supplementary Movie S4 (MP4)

Supplementary Movie S5 (MP4)

## Data availability

The data supporting the findings of this study are provided within this paper and its Supplementary materials.

## References

Barbier de Reuille, P., Routier-Kierzkowska, A. L., Kierzkowski, D., Bassel, G. W., Schüpbach, T., Tauriello, G., Bajpai, N., Strauss, S., Weber, A., Kiss, A., Burian, A., Hofhuis, H., Sapala, A., Lipowczan, M., Heimlicher, M. B., Robinson, S., Bayer, E. M., Basler, K., Koumoutsakos, P., Roeder, A. H.,…Smith, R. S., 2015). MorphoGraphX: A platform for quantifying morphogenesis in 4D. eLife, 4, 05864. 10.7554/eLife.05864.

Boruc, J., Mylle, E., Duda, M., De Clercq, R., Rombauts, S., Geelen, D., Hilson, P., Inzé, D., Van Damme, D., Russinova, E., 2010. Systematic localization of the Arabidopsis core cell cycle proteins reveals novel cell division complexes. Plant Physiol., 152, 553–565. 10.1104/pp.109.148643.

Brady, S. M., Orlando, D. A., Lee, J. Y., Wang, J. Y., Koch, J., Dinneny, J. R., Mace, D., Ohler, U., Benfey, P. N., 2007. A high-resolution root spatiotemporal map reveals dominant expression patterns. Science, 318, 801–806. 10.1126/science.1146265.

Chaves, M. M., Flexas, J., Pinheiro, C., 2009. Photosynthesis under drought and salt stress: regulation mechanisms from whole plant to cell. Ann. Bot., 103, 551–560. 10.1093/aob/mcn125.

Chun, H. J., Baek, D., Jin, B. J., Cho, H. M., Park, M. S., Lee, S. H., Lim, L. H., Cha, Y. J., Bae, D. W., Kim, S. T., Yun, D. J., & Kim, M. C., 2021. Microtubule dynamics plays a vital role in plant adaptation and tolerance to salt stress. Int. J. Mol. Sci., 22, 5957. 10.3390/ijms22115957.

Cifrová, P., Oulehlová, D., Kollárová, E., Martinek, J., Rosero, A., Žárský, V., Schwarzerová, K., Cvrčková, F., 2020. Division of labor between two actin nucleators – the formin FH1 and the ARP2/3 complex – in Arabidopsis epidermal cell morphogenesis. Front. Plant Sci., 11, 148. 10.3389/fpls.2020.00148.

Clough, S. J., Bent, A. F., 1998. Floral dip: a simplified method for Agrobacterium-mediated transformation of *Arabidopsis thaliana*. Plant J., 16, 735–743. 10.1046/j.1365-313x.1998.00343.x.

Cui, X., Zou, M., Li, J. (2023). Basally distributed actin array drives embryonic hypocotyl elongation during the seed-to-seedling transition in Arabidopsis. New Phytol., 240, 191–206. 10.1111/nph.19149.

Cvrčková F., 2019. From data to illustrations: common (free) tools for proper image data handling and processing. Methods Mol. Biol., 1992, 121–133. 10.1007/978-1-4939-9469-4_8.

Cvrčková, F., Ghosh, R., Kočová, H., 2024. Transmembrane formins as active cargoes of membrane trafficking. J. Exp. Bot., 75, 3668–3684. 10.1093/jxb/erae078.

Dabravolski, S. A., Isayenkov, S. V., 2023. The regulation of plant cell wall organisation under salt stress. Front. Plant Sci., 14, 1118313. 10.3389/fpls.2023.1118313.

Dai, L., Wang, B., Wang, T., Meyer, E. H., Kettel, V., Hoffmann, N., McFarlane, H. E., Li, S., Wu, X., Picard, K. L., Giavalisco, P., Persson, S., Zhang, Y. (2022). The TOR complex controls ATP levels to regulate actin cytoskeleton dynamics in Arabidopsis. Proc. Natl. Acad. Sci. USA, 119, e2122969119. 10.1073/pnas.2122969119.

Das, K. K., Mohapatra, A., George, A. P., Chavali, S., Witzel, K., Ramireddy, E., 2023. The proteome landscape of the root cap reveals a role for the jacalin-associated lectin JAL10 in the salt-induced endoplasmic reticulum stress pathway. Plant Commun., 4, 100726. 10.1016/j.xplc.2023.100726.

Deeks, M. J., Hussey, P. J., Davies, B., 2002. Formins: intermediates in signal-transduction cascades that affect cytoskeletal reorganization. Trends Plant Sci., 7, 492–498. 10.1016/s1360-1385(02)02341-5.

Deeks, M. J., Cvrčková, F., Machesky, L. M., Mikitová, V., Ketelaar, T., Žárský, V., Davies, B., Hussey, P. J., 2005. Arabidopsis group Ie formins localize to specific cell membrane domains, interact with actin-binding proteins and cause defects in cell expansion upon aberrant expression. New Phytol, 168, 529–540. 10.1111/j.1469-8137.2005.01582.x.

Diao, M., Ren, S., Wang, Q., Qian, L., Shen, J., Liu, Y., Huang, S., 2018. Arabidopsis formin 2 regulates cell-to-cell trafficking by capping and stabilizing actin filaments at plasmodesmata. eLife, 7, e36316. 10.7554/eLife.36316.

Dou, L., He, K., Higaki, T., Wang, X., Mao, T., 2018. Ethylene signaling modulates cortical microtubule reassembly in response to salt stress. Plant Physiol., 176, 2071–2081. 10.1104/pp.17.01124.

Duan, W. J., Liu, Z. H., Bai, J. F., Yuan, S. H., Li, Y. M., Lu, F. K., Zhang, T. B., Sun, J. H., Zhang, F. T., Zhao, C. P., Zhang, L. P. (2021). Comprehensive analysis of formin gene family highlights candidate genes related to pollen cytoskeleton and male fertility in wheat (*Triticum aestivum* L.). BMC Genomics, 22, 570. 10.1186/s12864-021-07878-7.

Engler, C., Kandzia, R., Marillonnet, S., 2008. A one pot, one step, precision cloning method with high throughput capability. PloS One, 3, e3647. 10.1371/journal.pone.0003647.

Fitz Gerald, J. N., Hui, P. S., Berger, F. (2009). Polycomb group-dependent imprinting of the actin regulator AtFH5 regulates morphogenesis in *Arabidopsis thaliana*. Development, 136, 3399–3404. 10.1242/dev.036921.

Fitzpatrick, M. 2014. Measuring cell fluorescence using ImageJ. https://theolb.readthedocs.io/en/latest/imaging/measuring-cell-fluorescence-using-imagej.html (accessed 4 November 2024),

Geldner, N., Anders, N., Wolters, H., Keicher, J., Kornberger, W., Muller, P., Delbarre, A., Ueda, T., Nakano, A., Jürgens, G., 2003. The Arabidopsis GNOM ARF-GEF mediates endosomal recycling, auxin transport, and auxin-dependent plant growth. Cell, 112, 219–230. 10.1016/s0092-8674(03)00003-5.

Gigli-Bisceglia, N., van Zelm, E., Huo, W., Lamers, J., Testerink, C., 2022. Arabidopsis root responses to salinity depend on pectin modification and cell wall sensing. Development, 149, dev200363. 10.1242/dev.200363.

Goh, T., Sakamoto, K., Wang, P., Kozono, S., Ueno, K., Miyashima, S., Toyokura, K., Fukaki, H., Kang, B. H., Nakajima, K.., 2022. Autophagy promotes organelle clearance and organized cell separation of living root cap cells in *Arabidopsis thaliana*. Development, 149, dev200593. 10.1242/dev.200593.

Grunt, M., Žárský, V., Cvrčková, F., 2008. Roots of angiosperm formins: the evolutionary history of plant FH2 domain-containing proteins. BMC Evol. Biol., 8, 115. 10.1186/1471-2148-8-115.

Ingouff, M., Fitz Gerald, J. N., Guérin, C., Robert, H., Sørensen, M. B., Van Damme, D., Geelen, D., Blanchoin, L., Berger, F., 2005. Plant formin AtFH5 is an evolutionarily conserved actin nucleator involved in cytokinesis. Nature Cell Biol., 7, 374–380. 10.1038/ncb1238.

Jaillais, Y., Fobis-Loisy, I., Miège, C., Gaude, T., 2008. Evidence for a sorting endosome in Arabidopsis root cells. Plant J., 53, 237–247. 10.1111/j.1365-313X.2007.03338.x.

Jiang, K., Moe-Lange, J., Hennet, L., Feldman, L. J., 2016. Salt stress affects the redox status of Arabidopsis root meristems. Front. Plant Sci., 7, 81. 10.3389/fpls.2016.00081.

Karimi, M., De Meyer, B., Hilson, P, 2005. Modular cloning in plant cells. Trends Plant Sci., 10, 103– 105. 10.1016/j.tplants.2005.01.008.

Kilian, J., Whitehead, D., Horak, J., Wanke, D., Weinl, S., Batistic, O., D’Angelo, C., Bornberg-Bauer, E., Kudla, J., Harter, K. (2007). The AtGenExpress global stress expression data set: protocols, evaluation and model data analysis of UV-B light, drought and cold stress responses. Plant J., 50, 347–363. 10.1111/j.1365-313X.2007.03052.x.

Klepikova, A. V., Kasianov, A. S., Gerasimov, E. S., Logacheva, M. D., Penin, A. A., 2016. A high resolution map of the *Arabidopsis thaliana* developmental transcriptome based on RNA-seq profiling. Plant J., 88, 1058–1070. 10.1111/tpj.13312.

Klionsky, D. J., Abdel-Aziz, A. K., Abdelfatah, S., Abdellatif, M., Abdoli, A., Abel, S., Abeliovich, H., Abildgaard, M. H., Abudu, Y. P., Acevedo-Arozena, A., Adamopoulos, I. E., Adeli, K., Adolph, T. E., Adornetto, A., Aflaki, E., Agam, G., Agarwal, A., Aggarwal, B. B., Agnello, M., Agostinis, P.,…Tong, C. K., 2021. Guidelines for the use and interpretation of assays for monitoring autophagy (4th edition). Autophagy, 17, 1–382. 10.1080/15548627.2020.1797280.

Knoblauch, M., Vendrell, M., de Leau, E., Paterlini, A., Knox, K., Ross-Elliot, T., Reinders, A., Brockman, S. A., Ward, J., Oparka, K., 2015. Multispectral phloem-mobile probes: properties and applications. Plant Physiol., 167, 1211–1220. 10.1104/pp.114.255414.

Knox K., 2019. Measuring phloem transport velocity in Arabidopsis seedlings using the fluorescent coumarin glucoside, esculin. Methods Mol. Biol., 2014, 195–201. 10.1007/978-1-4939-9562-2_16.

Kollárová, E., Baquero Forero, A., Stillerová, L., Přerostová, S., Cvrčková, F., 2020. Arabidopsis class II formins AtFH13 and AtFH14 can form heterodimers but exhibit distinct patterns of cellular localization. Int. J. Mol. Sci., 21, 348. 10.3390/ijms21010348.

Kollárová, E., Baquero Forero, A., Cvrčková, F., 2021. The *Arabidopsis thaliana* Class II formin FH13 modulates pollen tube growth. Front. Plant Sci., 12, 599961. 10.3389/fpls.2021.599961.

Kumar, S., Jeevaraj, T., Yunus, M. H., Chakraborty, S., Chakraborty, N., 2023. The plant cytoskeleton takes center stage in abiotic stress responses and resilience. Plant Cell Environ., 46, 5–22. 10.1111/pce.14450.

Kumar, M., Rani, K., 2023. Epigenomics in stress tolerance of plants under the climate change. Mol. Biol. Rep., 50, 6201–6216. 10.1007/s11033-023-08539-6.

Lan, Y., Liu, X., Fu, Y., Huang, S., 2018. Arabidopsis class I formins control membrane-originated actin polymerization at pollen tube tips. PLoS Genet., 14, e1007789. 10.1371/journal.pgen.1007789.

Lara-Mondragón, C. M., Dorchak, A., MacAlister, C. A., 2022. O-glycosylation of the extracellular domain of pollen class I formins modulates their plasma membrane mobility. J. Exp. Bot., 73, 3929– 3945. 10.1093/jxb/erac131.

Lei, Y., Lu, L., Liu, H. Y., Li, S., Xing, F., Chen, L. L., 2014. CRISPR-P: a web tool for synthetic single-guide RNA design of CRISPR-system in plants. Mol. Plant, 7, 1494–1496. 10.1093/mp/ssu044.

Li, C., Lu, H., Li, W., Yuan, M., Fu, Y., 2017. A ROP2-RIC1 pathway fine-tunes microtubule reorganization for salt tolerance in Arabidopsis. Plant Cell Environ., 40, 1127–1142. 10.1111/pce.12905.

Liu, S. G., Zhu, D. Z., Chen, G. H., Gao, X. Q., Zhang, X. S., 2012. Disrupted actin dynamics trigger an increment in the reactive oxygen species levels in the Arabidopsis root under salt stress. Plant Cell Rep., 31, 1219–1226. 10.1007/s00299-012-1242-z.

Liu, C., Zhang, Y., Ren, H., 2018. Actin polymerization mediated by AtFH5 directs the polarity establishment and vesicle trafficking for pollen germination in Arabidopsis. Mol. Plant, 11, 1389– 1399. 10.1016/j.molp.2018.09.004.

Liu, C., Zhang, Y., Ren, H., 2021. Profilin promotes formin-mediated actin filament assembly and vesicle transport during polarity formation in pollen. Plant Cell, 33, 1252–1267. 10.1093/plcell/koab027.

Liu, J., Zhang, W., Long, S., Zhao, C., 2021. Maintenance of cell wall integrity under high salinity. Int. J. Mol. Sci., 22, 3260. 10.3390/ijms22063260.

Liu, Z., Zou, L., Chen, C., Zhao, H., Yan, Y., Wang, C., Liu, X., 2019. iTRAQ-based quantitative proteomic analysis of salt stress in *Spica Prunellae*. Sci. Rep., 9, 9590. 10.1038/s41598-019-46043-9

Luo, L., Zhang, P., Zhu, R., Fu, J., Su, J., Zheng, J., Wang, Z., Wang, D., Gong, Q., 2017. Autophagy is rapidly induced by salt stress and is required for salt tolerance in Arabidopsis. Front.Plant Sci., 8, 1459. 10.3389/fpls.2017.01459.

Ma, H., Liu, M., 2019. The microtubule cytoskeleton acts as a sensor for stress response signaling in plants. Mol. Biol. Rep., 46, 5603–5608. 10.1007/s11033-019-04872-x.

Munns R., 2005. Genes and salt tolerance: bringing them together. New Phytol., 167, 645–663. 10.1111/j.1469-8137.2005.01487.x.

Munns, R., Millar, A. H., 2023. Seven plant capacities to adapt to abiotic stress. J. Exp. Bot., 74, 4308–4323. 10.1093/jxb/erad179.

Oulehlová, D., Kollárová, E., Cifrová, P., Pejchar, P., Žárský, V., Cvrčková, F., 2019. Arabidopsis class I formin FH1 relocates between membrane compartments during root cell ontogeny and associates with plasmodesmata. Plant Cell Physiol., 60, 1855–1870. 10.1093/pcp/pcz102.

Renna, L., Hanton, S. L., Stefano, G., Bortolotti, L., Misra, V.,Brandizzi, F., 2005. Identification and characterization of AtCASP, a plant transmembrane Golgi matrix protein. Plant Mol. Biol., 58, 109– 122. 10.1007/s11103-005-4618-4.

Rigal, A., Doyle, S. M., Robert, S., 2015. Live cell imaging of FM4-64, a tool for tracing the endocytic pathways in Arabidopsis root cells. Methods Mol. Biol., 1242, 93–103. 10.1007/978-1-4939-1902-4_9.

Ritzenthaler, C., Nebenführ, A., Movafeghi, A., Stussi-Garaud, C., Behnia, L., Pimpl, P., Staehelin, L. A., Robinson, D. G., 2002. Reevaluation of the effects of brefeldin A on plant cells using tobacco Bright Yellow 2 cells expressing Golgi-targeted green fluorescent protein and COPI antisera. Plant Cell, 14, 237–261. 10.1105/tpc.010237.

Rosero, A., Oulehlová, D., Stillerová, L., Schiebertová, P., Grunt, M., Žárský, V., Cvrčková, F., 2016. Arabidopsis FH1 formin affects cotyledon pavement cell shape by modulating cytoskeleton dynamics. Plant Cell Physiol., 57, 488–504. 10.1093/pcp/pcv209.

Roy, P. R., Tahjib-Ul-Arif, M., Polash, M. A. S., Hossen, M. Z., Hossain, M. A., 2019. Physiological mechanisms of exogenous calcium on alleviating salinity-induced stress in rice (*Oryza sativa* L.). Physiol. Mol. Biol. Plants, 25, 611–624. 10.1007/s12298-019-00654-8.

Ruan, H., Wang, T., Ren, H., Zhang, Y., 2023. AtFH5-labeled secretory vesicles-dependent calcium oscillation drives exocytosis and stepwise bulge during pollen germination. Cell Rep., 42, 113319. 10.1016/j.celrep.2023.113319.

Sengupta, S., Mangu, V., Sanchez, L., Bedre, R., Joshi, R., Rajasekaran, K., Baisakh, N., 2019. An actin-depolymerizing factor from the halophyte smooth cordgrass, Spartina alterniflora (SaADF2), is superior to its rice homolog (OsADF2) in conferring drought and salt tolerance when constitutively overexpressed in rice. Plant Biotech. J., 17, 188–205. 10.1111/pbi.12957.

Schindelin, J., Rueden, C. T., Hiner, M. C., Eliceiri, K. W. (2015). The ImageJ ecosystem: An open platform for biomedical image analysis. Mol. Reprod. Dev., 82, 518–529. 10.1002/mrd.22489.

Shabala, S., Demidchik, V., Shabala, L., Cuin, T. A., Smith, S. J., Miller, A. J., Davies, J. M., Newman, I. A., 2006. Extracellular Ca2+ ameliorates NaCl-induced K+ loss from Arabidopsis root and leaf cells by controlling plasma membrane K+-permeable channels. Plant Physiol., 141, 1653–1665. 10.1104/pp.106.082388.

Slovak, R., Göschl, C., Su, X., Shimotani, K., Shiina, T., Busch, W., 2014. A scalable open-source pipeline for large-scale root phenotyping of Arabidopsis. Plant Cell, 26, 2390–2403. 10.1105/tpc.114.124032.

Spitzer, M., Wildenhain, J., Rappsilber, J., Tyers, M., 2014. BoxPlotR: a web tool for generation of box plots. Nature Methods, 11, 121–122. 10.1038/nmeth.2811.

Stangroom, J., 2024. Social science statistics. https://www.socscistatistics.com/ (accessed 31 October 2024),

Steinhorst, L., He, G., Moore, L. K., Schültke, S., Schmitz-Thom, I., Cao, Y., Hashimoto, K., Andrés, Z., Piepenburg, K., Ragel, P., Behera, S., Almutairi, B. O., Batistič, O., Wyganowski, T., Köster, P., Edel, K. H., Zhang, C., Krebs, M., Jiang, C., Guo, Y.,…Kudla, J. (2022). A Ca^2+-^sensor switch for tolerance to elevated salt stress in Arabidopsis. Dev. Cell, 57, 2081–2094.e7. 10.1016/j.devcel.2022.08.001.

Takáč, T., Pechan, T., Šamajová, O., Ovečka, M., Richter, H., Eck, C., Niehaus, K., Šamaj, J., 2012. Wortmannin treatment induces changes in Arabidopsis root proteome and post-Golgi compartments. J. Proteome Res., 11, 3127–3142. 10.1021/pr201111n.

Tse, Y. C., Mo, B., Hillmer, S., Zhao, M., Lo, S. W., Robinson, D. G., Jiang, L., 2004. Identification of multivesicular bodies as prevacuolar compartments in Nicotiana tabacum BY-2 cells. Plant Cell, 16, 672–693. 10.1105/tpc.019703.

Ueda, T., Uemura, T., Sato, M. H., Nakano, A., 2004. Functional differentiation of endosomes in Arabidopsis cells. Plant J., 40, 783–789. 10.1111/j.1365-313X.2004.02249.x.

University of Toronto, 2024. The bio-analytic resources for plant biology. https://bar.utoronto.ca/ (accessed 31 October 2024).

van Zelm, E., Zhang, Y., Testerink, C., 2020. Salt tolerance mechanisms of plants. Annu. Rev. Plant Biol., 71, 403–433. 10.1146/annurev-arplant-050718-100005.

Vasavada, N., 2016. Online web statistical calculators. https://astatsa.com (accessed 31 October 2024),

Vicré, M., Santaella, C., Blanchet, S., Gateau, A., Driouich, A., 2005. Root border-like cells of Arabidopsis. Microscopical characterization and role in the interaction with rhizobacteria. Plant Physiol., 138, 998–1008. 10.1104/pp.104.051813.

Waadt, R., Seller, C. A., Hsu, P. K., Takahashi, Y., Munemasa, S., Schroeder, J. I., 2022. Plant hormone regulation of abiotic stress responses. Nat. Rev. Mol. Cell Biol., 23, 680–694. 10.1038/s41580-022-00479-6.

Wang, C., Li, J., Yuan, M., 2007. Salt tolerance requires cortical microtubule reorganization in Arabidopsis. Plant Cell Physiol., 48, 1534–1547. 10.1093/pcp/pcm123.

Wang, C., Zhang, L., Yuan, M., Ge, Y., Liu, Y., Fan, J., Ruan, Y., Cui, Z., Tong, S., Zhang, S., 2010. The microfilament cytoskeleton plays a vital role in salt and osmotic stress tolerance in Arabidopsis. Plant Biol., 12, 70–78. 10.1111/j.1438-8677.2009.00201.x.

Wang, L., Qiu, T., Yue, J., Guo, N., He, Y., Han, X., Wang, Q., Jia, P., Wang, H., Li, M., Wang, C., Wang, X., 2021. Arabidopsis ADF1 is regulated by MYB73 and is involved in response to salt stress affecting actin filament organization. Plant Cell Physiol., 62, 1387–1395. 10.1093/pcp/pcab081.

Wilkins, K. A., Matthus, E., Swarbreck, S. M., Davies, J. M. (2016). Calcium-mediated abiotic stress signaling in roots. Front. Plant Sci., 7, 1296. 10.3389/fpls.2016.01296.

Winter, D., Vinegar, B., Nahal, H., Ammar, R., Wilson, G. V., Provart, N. J., 2007. An “Electronic Fluorescent Pictograph” browser for exploring and analyzing large-scale biological data sets. PloS One, 2, e718. 10.1371/journal.pone.0000718.

Xing, H. L., Dong, L., Wang, Z. P., Zhang, H. Y., Han, C. Y., Liu, B., Wang, X. C., Chen, Q. J., 2014. A CRISPR/Cas9 toolkit for multiplex genome editing in plants. BMC Plant Biol., 14, 327. 10.1186/s12870-014-0327-y.

Xu, Y., Shen, J., Ruan, H., Qu, X., Li, Y., Wang, Y., Li, P., Yi, R., Ren, H., Zhang, Y., Huang, S. 2024. A RhoGAP controls apical actin polymerization by inhibiting formin in Arabidopsis pollen tubes. Curr. Biol., 34, 5040–5053. 10.1016/j.cub.2024.09.053.

Yang, J., An, B., Luo, H., He, C., Wang, Q., 2019. AtKATANIN1 modulates microtubule depolymerization and reorganization in response to salt stress in Arabidopsis Int. J. Mol. Sci., 21, 138. 10.3390/ijms21010138.

Ye, J., Zhang, W., Guo, Y., 2013. Arabidopsis SOS3 plays an important role in salt tolerance by mediating calcium-dependent microfilament reorganization. Plant Cell Rep., 32, 139–148. 10.1007/s00299-012-1348-3.

Zhang, H., Zhu, J., Gong, Z., Zhu, J. K., 2022. Abiotic stress responses in plants. Nat. Rev. Genet., 23, 104–119. 10.1038/s41576-021-00413-0.

Zhao, Y., Pan, Z., Zhang, Y., Qu, X., Zhang, Y., Yang, Y., Jiang, X., Huang, S., Yuan, M., Schumaker, S., Guo, Y., 2013. The actin-related protein 2/3 complex regulates mitochondrial-associated calcium signaling during salt stress in Arabidopsis. Plant Cell, 25, 4544–4559. 10.1105/tpc.113.117887.

Zwiewka, M., Nodzyński, T., Robert, S., Vanneste, S., Friml, J., 2015. Osmotic stress modulates the balance between exocytosis and clathrin-mediated endocytosis in *Arabidopsis thaliana*. Mol. Plant, 8, 1175–1187. 10.1016/j.molp.2015.03.007.

