## Supplementary Table, Figures and Legends for "The Arabidopsis Class I formin AtFH5 contributes to seedling resistance to salt stress"

### List of supplements in this file

**Table S1.** List and sequences of primers used in this study.

**Table S2**. Mendelian segregation of heterozygous AtFH5-GFP transgene in *fh5* mutant backgrounds.

**Figure S1**. The *fh5c* allele.

**Figure S2.** Expression pattern of AtFH5-GFP in additional *FH5* genotypes.

**Figure S3.** Association of AtFH5-GFP labelled bodies with the Golgi apparatus.

**Figure S4.** Co-localization of AtFH5-GFP with late endosomes.

**Figure S5.** Effect of sucrose on AtFH5-GFP expression and localization.

**Figure S6.** Mannitol treatment inhibits root growth but does not cause seedling lethality.

**Figure S7**. NaCl toxicity in both WT and *atfh5* seedlings is alleviated by $CaCl_2$ treatment.

### Additional supplementary files

**Movie S1.** SDCM Z-stack of pAtFH5::AtFH5-GFP transgenic root cap. Scale = 10µm.

**Movie S2.** Time lapse recording of AtFH5-GFP localization in root cap stained for 5 min by FM4-64. Scale = 10µm

**Movie S3.** Time lapse recording from a transiently transformed *N. benthamiana* leaf epidermis pavement cell co-expressing AtFH5-GFP and red fluorescent protein-labelled sialyltransferase (ST-RFP). Scale = 10µm.

**Movie S4.** Comparison of SDCM Z-stacks of pAtFH5::AtFH5-GFP root caps after mock treatment with DMSO (top) and WM. Scale = 10µm.

**Movie S5.** Time lapse recording of AtFH5-GFP and ARA6-RFP co-expression in epidermal cells of *N. benthamiana*. Scale = 10µm.

**Table S1.** List and sequences of primers used in this study.

| Primer name | Primer sequence 5´- 3´ | Purpose |
|---|---|---|
| DT1-BsF-FH5 | ATATATGGTCTCGATTGCCCTCCTAAGCGTAGTCGGGTT | AtFH5 gRNA cloning (with DT2-BsR-FH5) |
| DT2-BsR-FH5 | ATTATTGGTCTCGAAACCCGACTACGCTTAGGAGGGCAA | AtFH5 gRNA cloning (with DT2-BsF-FH5) |
| FH5seq_F | CAAAGGATAGTAGGGAAC | amplification and sequencing of the *fh5c* allele (with FH5seq_R) |
| FH5seq_R | CTCCACTTTCTCACTGT | amplification and sequencing of the *fh5c* allele (with FH5seq_F) |
| CAS9 fw | CTCGACTCACGGATGAACACTAA | detection of Cas9 cassette |
| CAS9 rv | AAATCTGCTCAATGATCTCGTCGA | detection of Cas9 cassette |
| AtFH5_full_for_gw | GGGGACAAGTTTGTACAAAAAAGCAGGCTTAATGGTTGGAATGATTCGAGGAGG | cloning of AtFH5 genomic sequence |
| AtFH5_full_rev_gw | GGGGACCACTTTGTACAAGAAAGCTGGGTAGTCTGAATCTGAACTAGACTGATCCAC | cloning of AtFH5 genomic sequence |
| AtFH5_for | ATGGTTGGAATGATTCGAGGAGG | PCR of AtFH5 before using GW primers |
| AtFH5_rev | GTCTGAATCTGAACTAGACTGATCCAC | PCR of AtFH5 before using GW primers |
| pFH5_rev_GW | GGGGACAACTTTGTATAGAAAAGTTGCTATCTTTTGCTTCTCATTTCAT | cloning of the AtFH5 promoter |
| pFH5_for_GW | GGGGACTGCTTTTTTGTACAAACTTGCGAGAGAAATTGAGACGAATG | cloning of the AtFH5 promoter |
| gFH5sek1 | GCAAAATAGTTTGGATTCGCCATTTG | gFH5 sequencing |
| gFH5sek2 | TGGTGGTTCGGTCAAAGGAG | gFH5 sequencing |
| gFH5sek3 | CAATTCTTTTGCGTGCTTTGAATGC | gFH5 sequencing |
| LP_SALK_152090 | TTTTCGATCAGGGTTGTTGAG | detection of T-DNA insertional allele *fh5-3* |
| RP_SALK_152090 | AAGAGCTCCAGATACTTGGGG | detection of T-DNA insertional allele *fh5-3* |

**Table S2**. Mendelian segregation of heterozygous pAtFH5::AtFH5-GFP transgene in *fh5* mutant backgrounds.

| Line | Signal | | No signal | | Total No. | $X^2$ | P-value |
|---|---|---|---|---|---|---|---|
| | No. | % | No. | % | | | |
| AtFH5-GFP-/+ in *fh5-3*-/- | 103 | 82 | 23 | 18 | 126 | 1.449 | 0.229 |
| AtFH5-GFP-/+ in *fh5c*-/- (9) | 100 | 77 | 30 | 23 | 130 | 0.110 | 0.741 |
| AtFH5-GFP-/+ in *fh5c*-/- (10) | 100 | 76 | 31 | 24 | 131 | 0.027 | 0.869 |
| Expected | | 75 | | 25 | | | |

For the *fh5c* background, data from two independent transformant lines (9) and (10) are shown.

A

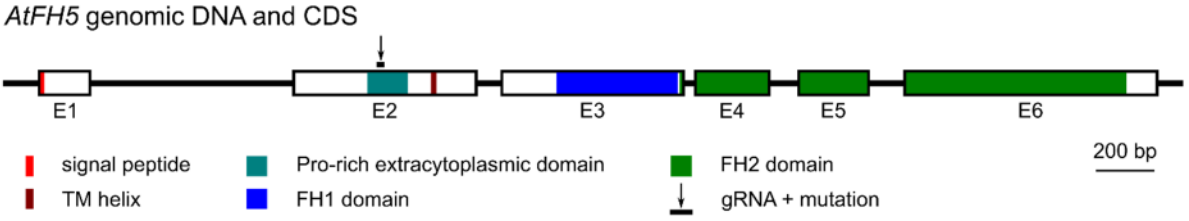


B

```
WT    CCTTCCCCTTCACCTTCCCGCCCTCCTAAGCGTAG-TCGGGGGCCTCCGCGCCCACCT
fh5c  CCTTCCCCTTCACCTTCCCGCCCTCCTAAGCGTAGTTCGGGGGCCTCCGCGCCCACCT
```

C

```
WT    MVGMIRGGMG DQNWSRLVFW LILFSGLLVI TLEENPEKDE IFLSQFMAPS TGQVNEHMEE TSWAQRCWQD SDCVKEAVAE
fh5c  MVGMIRGGMG DQNWSRLVFW LILFSGLLVI TLEENPEKDE IFLSQFMAPS TGQVNEHMEE TSWAQRCWQD SDCVKEAVAE

WT    FNLCFPGSKD SRELFGLNHT NLKQTLLDCI QEKGKLNGHN PKYLELLSSM LDIPRRNLAT KPGSSPSPSP SRPPKRSRGP
fh5c  FNLCFPGSKD SRELFGLNHT NLKQTLLDCI QEKGKLNGHN PKYLELLSSM LDIPRRNLAT KPGSSPSPSP SRPPKRSSGA

WT    PRPPTRPKSP PPRKSSFPPS RSPPPPPAKK NASKNSTSAP VSPAKKKEDH EKTIIIAVVV TAVSTFLLAA LFFLCCSRVC
fh5c  SAPTYSTKIP TSSEIIFSTI KISPSSSC*

WT    GNGSGGRKND ERPLLSLSSS DYSVGSSINY GGSVKGDKQG HQSFNIYSNQ GKMSSFDGSN SDTSDSLEER LSHEGLRNNS
      ITNHGLPPLK PPPGRTASVL SGKSFSGKVE PLPPEPPKFL KVSSKKASAP PPPVPAPQMP SSAGPPRPPP PAPPPGSGGP
      KPPPPPGPKG PRPPPPMSLG PKAPRPPSGP ADALDDDAPK TKLKPFFWDK VQANPEHSMV WNDIRSGSFQ FNEEMIESLF
      GYAAADKNKN DKKGSSGQAA LPQFVQILEP KKGQNLSILL RALNATTEEV CDALREGNEL PVEFIQTLLK MAPTPEEELK
      LRLYCGEIAQ LGSAERFLKA VVDIPFAFKR LEALLFMCTL HEEMAFVKES FQKLEVACKE LRGSRLFLKL LEAVLKTGNR
      MNDGTFRGGA QAFKLDTLLK LADVKGTDGK TTLLHFVVQE IIRTEGVRAA RTIRESQSFS SVKTEDLLVE ETSEESEENY
      RNLGLEKVSG LSSELEHVKK SANIDADGLT GTVLKMGHAL SKARDFVNSE MKSSGEESGF REALEDFIQN AEGSIMSILE
      EEKRIMALVK STGDYFHGKA GKDEGLRLFV IVRDFLIILD KSCKEVREAR GRPVRMARKQ GSTASASSET PRQTPSLDPR
      QKLFPAITER RVDQSSSDSD *
```

signal peptide
TM helix
Pro-rich extracytoplasmic domain
FH1 domain
FH2 domain
**out of frame segment**

**Figure S1**. **The *fh5c* allele.** (A) Map of the *AtFH5* locus with position of the guide RNA and the point mutation in the *fh5c* allele. The coding exons (E1 - E6) are coloured to reflect the domain structure of the AtFH5 protein. (B) Alignment of the WT and mutant genomic sequences. The segment corresponding to guide RNA is shown in bold, the inserted base in red. (C) Comparison of the protein sequence of WT AtFH5 and the fragment encoded by *fh5c*.

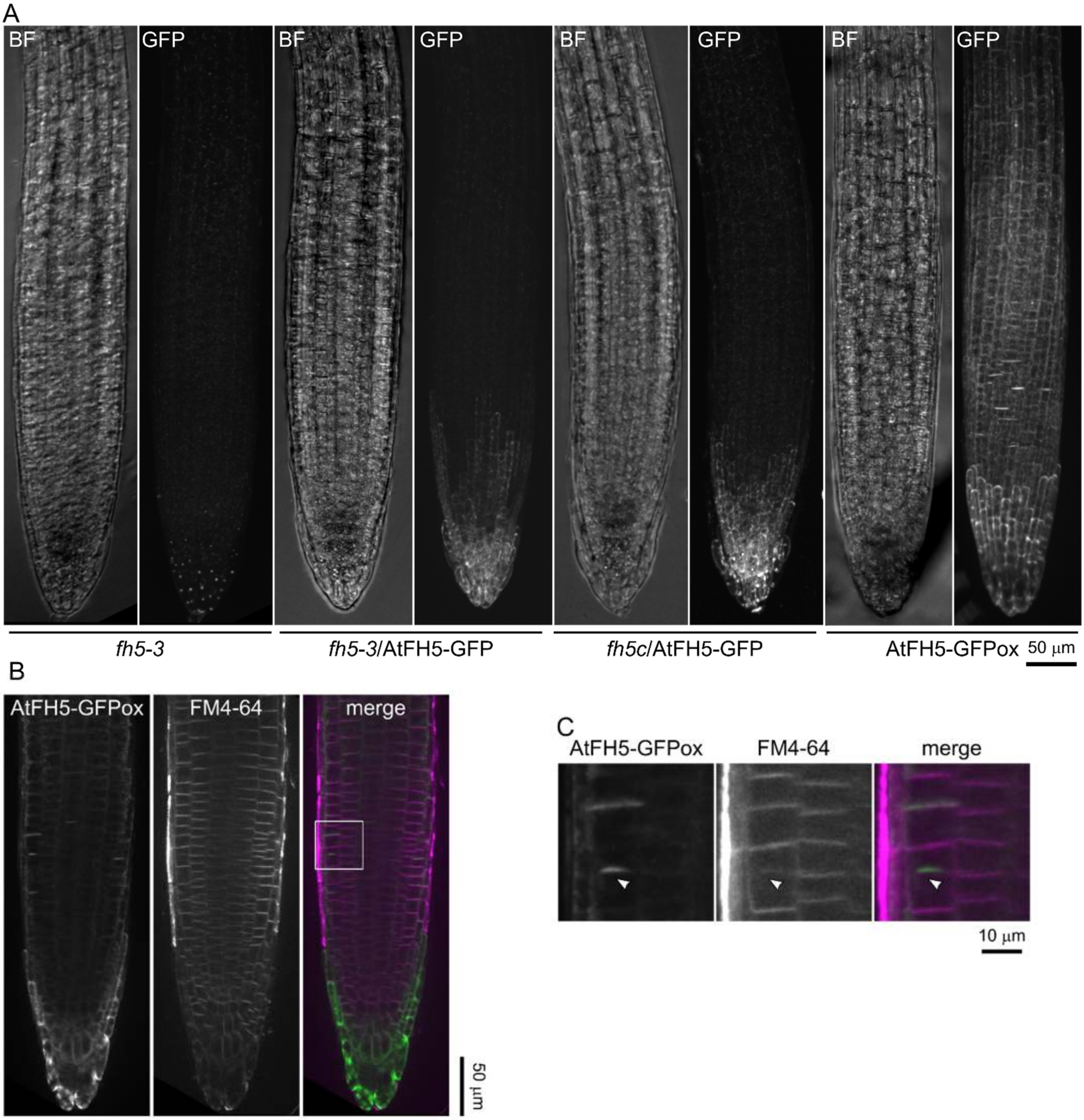


**Figure S2. Expression pattern of AtFH5-GFP in additional *FH5* genotypes.** (A) Z-projections of SDCM Z-stacks. Single channel images of bright field (BF) and GFP channels are shown. (B) A longitudinal SDCM optical section of an overexpressing AtFH5-GFPox root stained by FM4-64. Single channel images are shown in grayscale, in the merged display channels are represented by green (GFP) and magenta (FM4-64). (C) Close-up images of the marked area from (B) showing the localization of the developing meristematic cell plate (arrowhead).

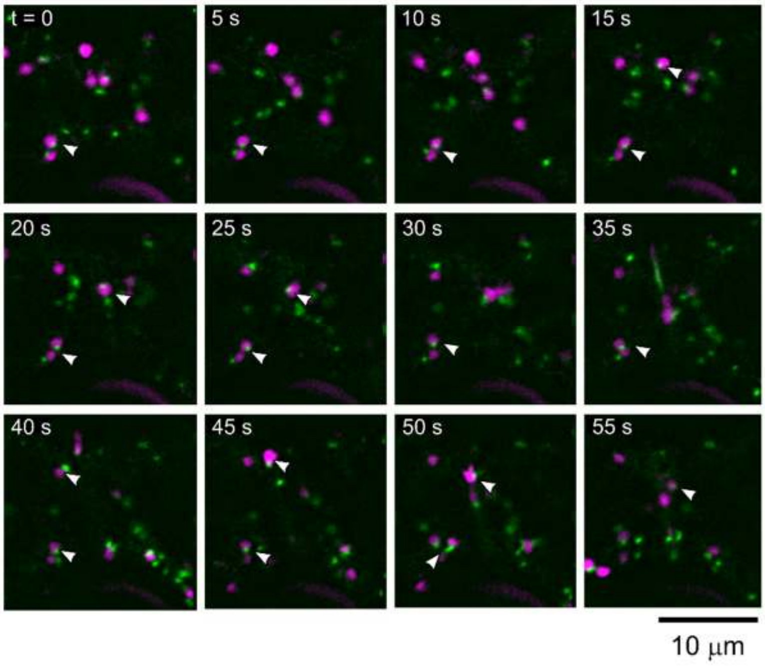


**Figure S3. Association of AtFH5-GFP labelled bodies with the Golgi apparatus.** Time lapse images of transiently transformed *N. benthamiana* leaf epidermis pavement cell co-expressing AtFH5-GFP (green) and red fluorescent protein-labelled sialyl transferase (ST-RFP, magenta). Arrowheads show mobile AtFH5-carrying particles traveling alongside Golgi bodies.

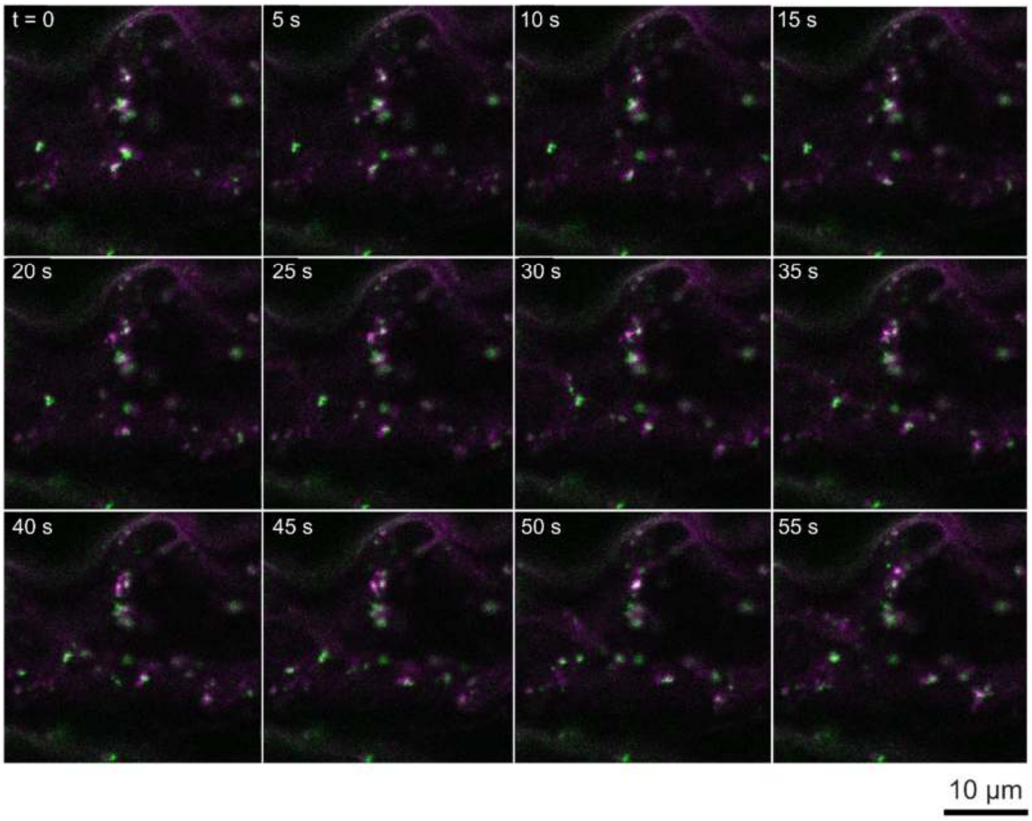


**Figure S4. Co-localization of AtFH5-GFP with late endosomes.** Time lapse images of transiently transformed *N. benthamiana* leaf epidermis pavement cell co-expressing AtFH5-GFP (green) and red fluorescent protein-labelled ARA6 GTPase (magenta).

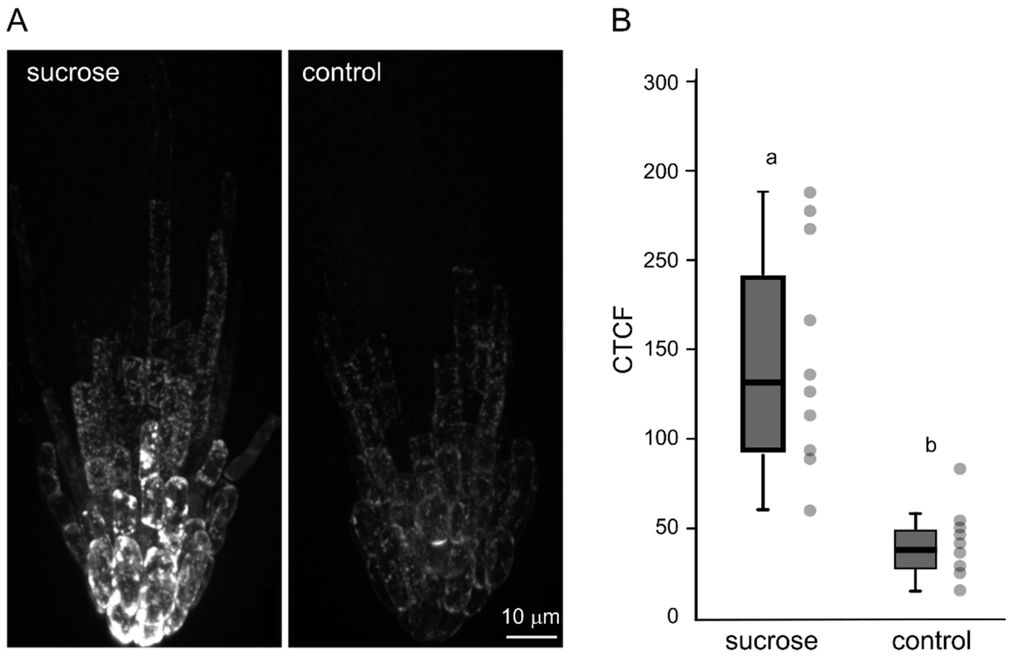


**Figure S5. Effect of sucrose on AtFH5-GFP expression and localization.** 3D reconstruction of a SDCM Z-stacks of a transgenic seedling root cap after cultivation with and without 1 % sucrose. Corrected total cell fluorescence (CTCF) measurement of FH5-GFP signal in border-like cells after cultivation with and without 1 % sucrose.

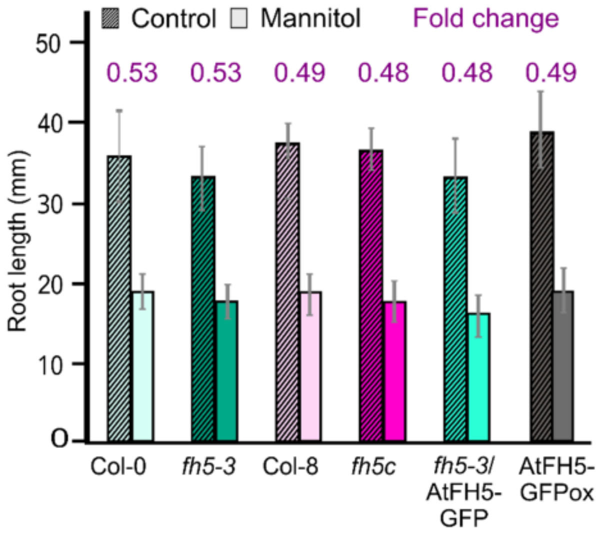


**Figure S6. Mannitol treatment inhibits root growth but does not cause seedling lethality**. Primary root length of seedlings of the indicated genotypes 5 days after transfer to a new standard medium plate or to medium containing 300 mM mannitol. Data presented are means ± SD, n ≥ 20 per genotype. Although some of the between-genotype differences in growth rate were statistically significant in any given experiment, this was not well reproducible between independent biological repeats. However, the extent of inhibition is similar for all genotypes (numbers denote fold change, i.e. the ratio between average root length of treated vs control seedlings).

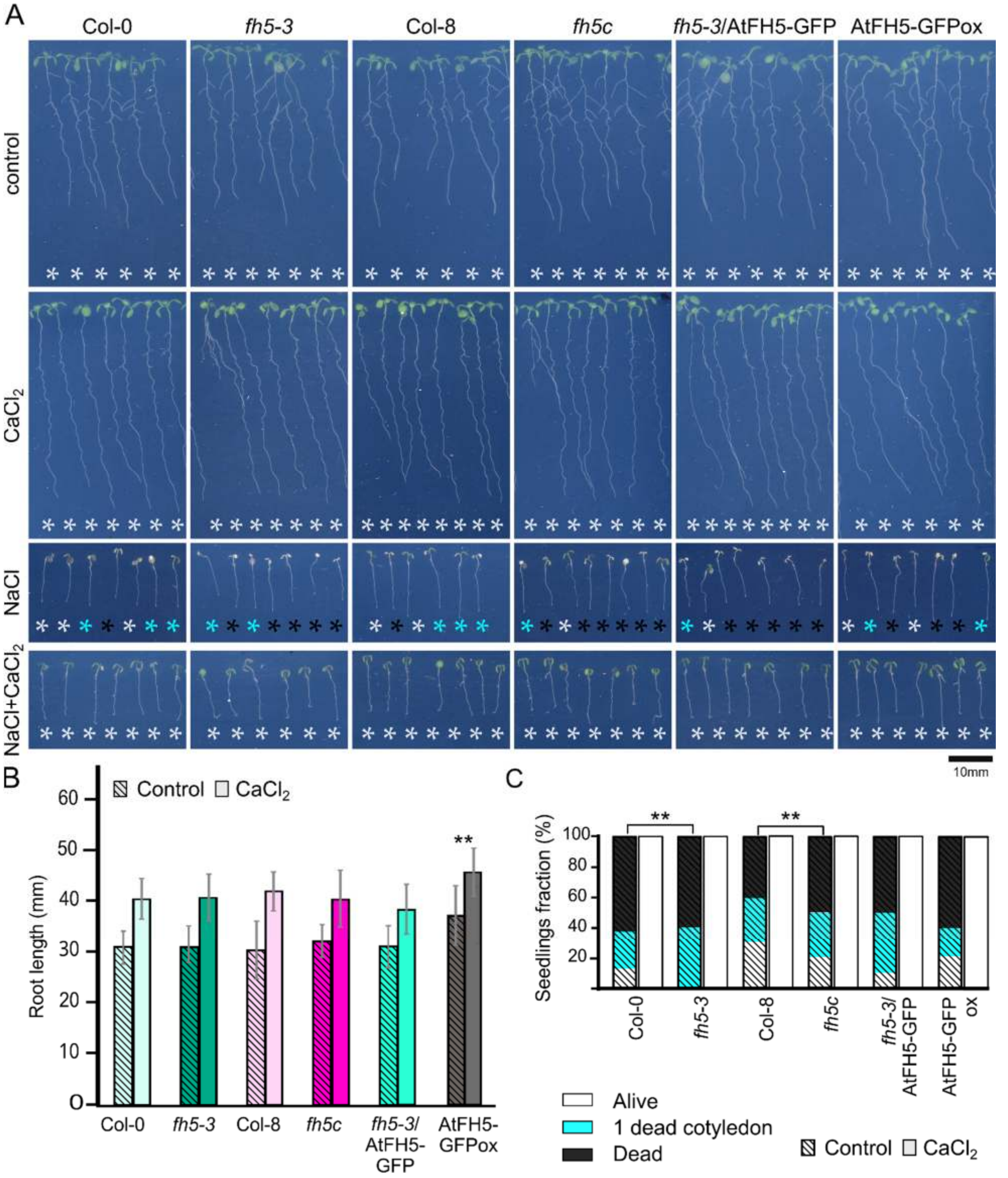


**Figure S7. NaCl toxicity in both WT and *atfh5* seedlings is alleviated by $CaCl_2$ treatment.** (A) Representative seedlings after 5 days at 0 and 250 mM NaCl, 10 mM $CaCl_2$ or both salts simultaneously. (B) Primary root length of seedlings of the indicated genotypes five days after transfer to a new standard medium or to medium containing 10 mM $CaCl_2$. Between-genotype differences are not statistically significant except for the overexpression line (one-way ANOVA, $p < 0.01$). Data shown are means ± SD, $n \geq 30$ per genotype. (C) Fraction of surviving or dead seedlings exhibiting various extent of damage after 5 days after transfer to media with either NaCl alone or NaCl and $CaCl_2$. Black – bleached seedlings, cyan – one bleached cotyledon, white – surviving seedlings. Asterisks denote significant differences ($p < 0.01$). Presence of $CaCl_2$ is indicated by cross-hatching in (B) and (C).
